# Comparative transcriptome profiling of the human and mouse dorsal root ganglia: An RNA-seq-based resource for pain and sensory neuroscience research

**DOI:** 10.1101/165431

**Authors:** Pradipta Ray, Andrew Torck, Lilyana Quigley, Andi Wangzhou, Matthew Neiman, Chandranshu Rao, Tiffany Lam, Ji-Young Kim, Tae Hoon Kim, Michael Q. Zhang, Gregory Dussor, Theodore J. Price

**Affiliations:** The University of Texas at Dallas, School of Behavioral and Brain Sciences; The University of Texas at Dallas, Department of Biological Sciences

## Abstract

Molecular neurobiological insight into human nervous tissues is needed to generate next generation therapeutics for neurological disorders like chronic pain. We obtained human Dorsal Root Ganglia (DRG) samples from organ donors and performed RNA-sequencing (RNA-seq) to study the human DRG (hDRG) transcriptional landscape, systematically comparing it with publicly available data from a variety of human and orthologous mouse tissues, including mouse DRG (mDRG). We characterized the hDRG transcriptional profile in terms of tissue-restricted gene co-expression patterns and putative transcriptional regulators, and formulated an information-theoretic framework to quantify DRG enrichment. Our analyses reveal an hDRG-enriched protein-coding gene set (~140), some of which have not been described in the context of DRG or pain signaling. A majority of these show conserved enrichment in mDRG, and were mined for known drug - gene product interactions. Comparison of hDRG and tibial nerve transcriptomes suggest pervasive mRNA transport of sensory neuronal genes to axons in adult hDRG, with potential implications for mechanistic insight into chronic pain in patients. Relevant gene families and pathways were also analyzed, including transcription factors (TFs), g-protein coupled receptors (GCPRs) and ion channels. We present our work as an online, searchable repository (http://www.utdallas.edu/bbs/painneurosciencelab/DRGtranscriptome), creating a valuable resource for the community. Our analyses provide insight into DRG biology for guiding development of novel therapeutics, and a blueprint for cross-species transcriptomic analyses.

**Summary:** We generated RNA sequencing data from human DRG samples and comprehensively compared this transcriptome to other human tissues and a matching panel of mouse tissues. Our analysis uncovered functionally enriched genes in the human and mouse DRG with important implications for understanding sensory biology and pain drug discovery.

## 1. Introduction

The dorsal root ganglia (DRG) is a primary sensory tissue in vertebrate nervous systems, delivering sensory signals from the body to the central nervous system (CNS) via pseudounipolar neurons. The DRG is composed of several specialized cell types including proprioceptive, low-threshold and damage-sensing nociceptive sensory neurons (nociceptors) as well as Schwann cells, fibroblasts and satellite glial cells (SGCs). With the advent of high-throughput RNA-seq, there has been a concerted effort on generating whole transcriptome snapshots of DRG and other tissues with recent emphasis on single cell RNA-seq (scRNA-seq) studies. Thus far, efforts have largely been directed toward mouse DRG (mDRG) [54] and rat DRG [46], with some focusing on single neuron [14; 51; 107] or specific neuronal subpopulations [29; 37; 102] highlighting nociceptors. Such studies are informative because they focus on profiling a key neuronal subpopulation in the context of detecting nociceptive signals and generation of pain, enhancing our understanding of the molecular biology and diversity of these neurons. The hDRG in general, and nociceptors in particular, are widely viewed as tissue and cell types with possibilities for discovering biological targets for analgesic drugs, leading to development of therapeutics for both acute and chronic pain [66; 92] that do not involve the CNS and avoid resulting complications [4; 19]. Studies identifying transcriptional [20; 32; 116; 119] (including scRNA-seq efforts [35]) and proteomic [89] changes in preclinical pain models have been undertaken, but do not account for evolutionary differences in DRG transcriptomes between rodent models and humans. Such preclinical models can potentially suffer from translational issues as they move toward the clinic [94]. Studies characterizing transcriptome profiles of human nociceptors and DRG (or related tissues and cell types *in vivo* or *in vitro*) present opportunities for advancing understanding of basic pain mechanisms and therapeutic targets but a limited number of such *in vitro* [111] or *in vivo* [25; 91; 98; 111] studies have been performed. The SymAtlas [98] project provided the first *in vivo*, high-throughput (but incomplete) characterization of the hDRG transcriptome using microarrays. Recent *in vivo* studies [25; 91] perform RNA-seq, but have mainly focused on specific gene families or diseases, without systematic comparison with preclinical models.

We performed RNA-seq on female, human organ donor L2 DRG tissues. Comprehensive analyses of the transcriptional landscape with respect to other human tissues identified gene co-expression modules and hDRG-enriched genes in pertinent gene families and signaling pathways. As a starting point for therapeutic target identification, we identified the set of genes with conserved DRG enrichment in human and mouse, and mined drug databases to catalog interactions of known drugs with products of these genes. This effort is the first to present whole transcriptome gene abundances for hDRGs and mDRGs, contrasted with relevant human and mouse tissues via our online repository (www.utdallas.edu/bbs/painneurosciencelab/DRGtranscriptome).

## 2. Methods

We performed RNA-seq for hDRG and integratively analyzed the transcriptome by comparing it with publicly available human and mouse RNA-seq datasets (sources shown in Table 1, and the workflow for the project shown in Figure 1). We also used publicly available human microarray data to contrast whole human DRG and Trigeminal Ganglia (TG) (GEO PRJNA87249) [98], along with two datasets representative of its component cell types: cultured primary Normal Human Schwann Cells (NHSCs) from peripheral nerves (GEO GSE14038) [62] and cultured fibroblasts (FIBRO) from the skin (GEO dataset GSE21899). We were unable to find a dataset for purified satellite glial cells (SGCs) in any database. We further integrated published quantification from mDRG scRNA-seq analyses [107] (lumbar DRG, GEO GSE59739) to putatively identify cell-type specific expression in DRG enriched genes. For ease of discourse, human gene names (in upper case) are used in the text to refer to both human genes and its orthologs, with the species being apparent from context. Mouse gene names and human protein names are capitalized, with human protein names being additionally italicized, when used in the paper. At the outset, it is important to clarify that our analyzed tissue panel only queries a finite set of adult, healthy tissue; and the notions of ‘DRG enriched’ and ‘DRG specific’ gene expression is with respect to this set of reference tissues that provide a relevant test-bench for asking questions about mammalian DRG.

**Figure 1.**
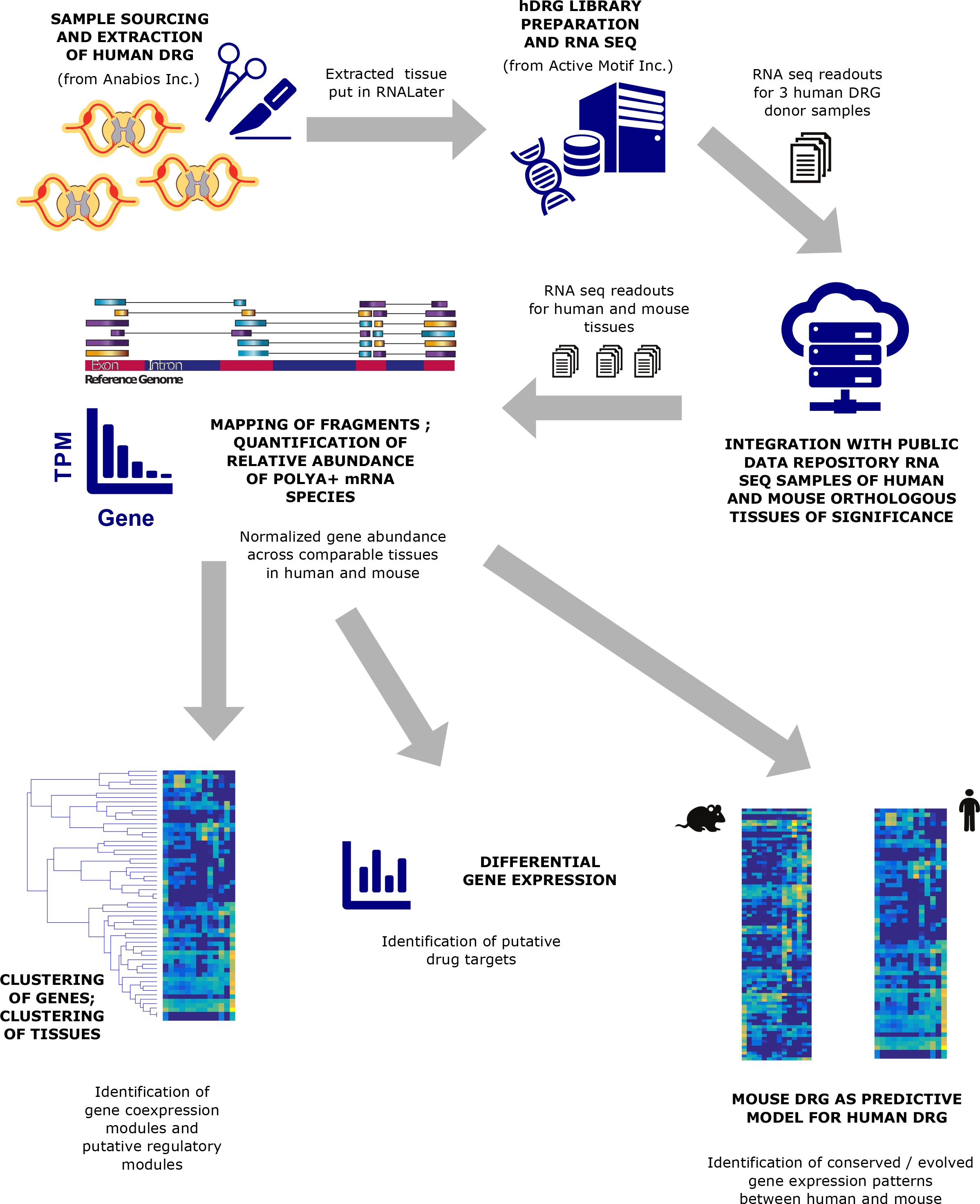
Schema of our study for generation and analysis of mRNA abundance profiles. The broad characterization of our work showing mRNA sequencing, RNA-seq mapping and quantification, gene expression normalization across samples and genes, and characterization of transcript profiles in terms of their tissue specificity, expression patterns across tissues, and conservation of expression pattern between human and preclinical mouse model is shown.

### 2.1 Human DRG preparation and RNA-seq

Tissue was sourced from Anabios, Inc (San Diego, CA). For this study, L2 lumbar DRGs were removed from 3 female consented organ donors prior to cross clamp and stored immediately in RNALater (Ambion, Austin, TX). RNA-seq was performed on the whole tissue by Active Motif, Inc (San Diego, CA). An RNEasy Qiagen kit was used for total RNA isolation, and DNA was removed using DNase-I digestion. The extracted total RNA was then used with an Illumina Truseq RNA sample preparation v2 kit to generate polyA+ RNA libraries for sequencing. 50 million paired-end 75 bp reads were sequenced from each sample on the Illumina platform. The RNA-seq datasets generated were contrasted with publicly available RNA-seq datasets in other relevant human tissues, and their orthologous tissues in the house mouse (*M. musculus*). For mDRG RNA-seq datasets, we used publicly available RNA-seq data from the C57BL/6 mouse strain, the inbred model strain used to sequence the reference mouse genome [13].

### 2.2 Quantification of gene abundances from RNA-seq experimental data

*Mapping of sequenced reads and quantification of gene abundances*: Sequenced FASTQ files generated by our RNA-seq experiments or downloaded from public databases were mapped to the human and mouse reference transcriptome for the purposes of quantifying relative gene abundance. Reference human (NCBI hg19) and mouse (NCBI mm10) genomes [21], and reference human (Gencode v14) and mouse (Gencode vM4) transcriptomes [33] were used for mapping sequenced reads. The Tophat / Cufflinks pipeline was used for analyzing the RNA-seq datasets [105]. The Burroughs-Wheeler transform based RNA-seq mapping tool Tophat [104] (v2.0.13) was used for mapping the sequencing reads with the following command-line parameters: *`tophat2 -o <output_path> -p 8 --transcriptome-index <reference_transcriptome> <reference_genome> <leftreads.fastq> <rightreads.fastq>`*. The gene abundance quantification tool Cuffdiff [103] (v2.2.1) was used to estimate relative abundances of genes in the reference transcriptome with the following command-line parameters: *`cuffdiff -o <output_file> --library-norm-method classic-fpkm -p 8 <reference_transcriptome>`*. Relative abundances were calculated for genes with respect to the reference library, based on Gencode coding genes, lincRNAs, pseudogenes and other noncoding genes with evidence for polyA site or signal [121]. Absolute abundance quantification of genes or transcripts require calibrating RNA-seq based on exogenous spike-ins with known copy numbers [78] or other methods is rarely performed in practice, and typically require additional amounts of input material, or attachment of unique molecular barcodes to individual transcripts to correct amplification bias in traditional RNA-seq [96], making them unfeasible or intractable for our samples with limited, varying amounts of tissue sourced from human donors.

*Design decisions for RNA-seq analyses*: While mapping RNA-seq datasets using Tophat, the number of mismatches allowed per alignment segment was set to the default (two), with all of our analyzed datasets, including the three hDRG samples we sequenced, having acceptable mapping rates for post-mortem samples (> 70%) [87]. For Tophat, all datasets were assumed to be generated from strand-agnostic libraries for the purposes of mapping. While some analyzed libraries were actually strand-specific, such an approach allows us to avoid the inherently different nature of strand specific and non-specific libraries, and minimizes the chances that downstream differential expression analysis will yield artifacts. Since the reference transcriptomes for human and mouse are well annotated by the GENCODE project, *de novo* assembly was not performed - instead, all concordantly mapped read pairs (fragments) mapping to each known transcript were used by Cuffdiff to quantify relative abundance. Finally, we additionally performed a second round of *in silico* library selection limiting analysis to only genes with known or predicted polyA+ transcripts (detailed in Section 2.3 under TPM calculations), to minimize the effect of whether total RNA or polyA+ selection was used for library construction in the different analyzed samples.

### 2.3 Sample statistics

#### Sample statistics for characterizing gene abundances in individual tissues for each species

The relative abundance of a transcript was calculated based on the number of fragments (paired-end reads) that map to it, by calculating Fragments per Kilobase per Million Mapped Fragments (FPKM) as follows:

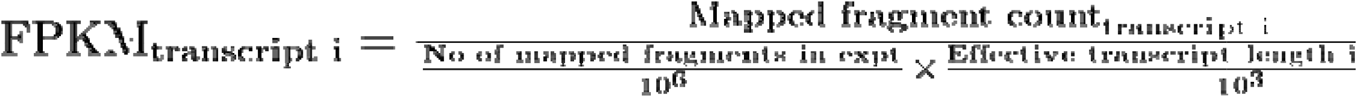

FPKMs for genes were calculated by summing the FPKMs for individual transcripts corresponding to the gene. Cuffdiff was configured to use all replicates to estimate a mean FPKM for each gene. Our analysis ultimately required us to compare across tissues and species for many different RNA-seq experiments, performed across multiple laboratories utilizing different library preparations and sequencing depth. Hence, we decided to transform the FPKMs to another commonly used measure for quantifying relative abundance: Transcripts per million (TPM) [79]. To calculate TPMs, FPKMs are re-normalized with respect to the sum of FPKMs of the reference library of transcripts to generate a relative abundance score scaled to parts per million.

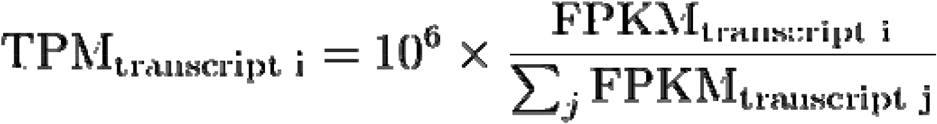

We only chose to quantify relative abundance with respect to transcripts that were polyA+, since most datasets in our analysis used a polyA selection step. LincRNAs, pseudogenes and other noncoding genes with evidence of polyA signal or site (based on APASdb [121]) were retained in the reference library, while remaining noncoding genes (including ribosomal rRNA genes) were not evaluated. By only quantifying the relative abundance of genes with respect to the set of common genes that are minimally targeted by most RNA-seq protocols, we effectively performed a second round of library selection *in silico*. Finally, in order to scale the distribution of TPMs in each tissue comparably, the upper quartile (75^th^ percentile) TPM was calculated for each sample and values scaled with respect to it to calculate upper quartile-normalized TPMs (uqTPM), based on previous approaches in the literature [28]. A complete list of evaluated genes, with relative abundances in TPM and uqTPM across all analyzed tissues, is available in Supplementary Files 1 (human) and 2 (mouse). Empirical density functions for uqTPMs for different tissues in both human and mouse are presented in Supplementary Fig. 1.

We used upper quartile normalization to scale TPM measures such that the bimodal distribution of relative abundances were comparable across both tissues and species. Based on the bimodal empirical density functions of the uqTPMs in different tissues for both species, we categorized genes as ““expressed” versus “undetected or lowly expressed”, when the mean abundance across replicates was >= 0.75 uqTPM. Additionally, this allows for simplification of downstream analyses like co-expression module detection, identifying regulatory networks or gene set enrichment analysis [11].

#### Sample statistics for characterizing tissue specificity of genes in individual species

A tissue-specific transcriptomic signature inevitably depends upon a set of tissue-restricted genes, potentially contributing to tissue-specific functionality or phenotype. The information theoretic measure of Shannon’s entropy [95] has been historically used in various domains as a framework for assessing diversity in a population [34], including the identification of tissue-restricted genes in transcriptomics [117]. For each gene, the uqTPM levels were first normalized to 1, generating a probability distribution 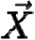 with X_i_ corresponding to the gene’s normalized abundance and t_i_ corresponding to its uqTPM in the *i* th tissue. Based on this distribution, Shannon’s entropy [95] 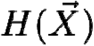 was calculated to identify tissue-restricted genes that are expressed in the DRG:

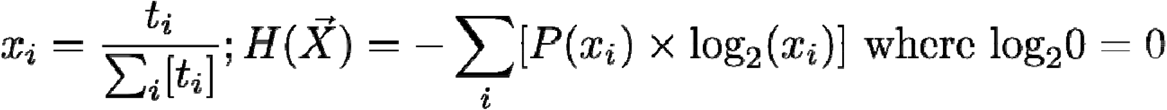

Since the value of Shannon’s entropy can vary between 0 and log_2_(n) where n = information length (number of tissues in our case), we use Shannon’s normalized measure of entropy which ranges between 0 (highest tissue specificity) and 1 (lowest tissue specificity): 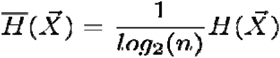

We defined two additional formulations derived from this framework to characterize tissue-restricted genes of interest. We profiled nervous-system enriched genes among the set of tissue-restricted genes we characterized in this work, since these are potential targets or excluded targets for drugs which do or do not cross the blood brain barrier (BBB). We calculated a neural proportion score by summing the proportion P(x_i_) calculated for Shannon’s entropy across all neural tissues in our tissue panel. We also calculated a score for DRG enrichment that would take into account the magnitude of relative abundance values across different tissues. For each gene, we also calculated a DRG enrichment score to quantify enrichment in the DRG with respect to other analyzed tissues, ranging from 0 (undetected in the DRG or uniformly expressed across tissues) to 1 (only expressed in the DRG), defined as 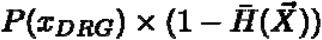. Empirical density functions for the neural proportion and DRG enrichment scores for both human and mouse are presented in Supplementary Fig. 2. The degree of neural or DRG enrichment for a gene can thus be estimated by contrasting the enrichment score with the overall distribution of the score across all genes.

#### Sample statistics for characterizing conservation of gene expression patterns in human and mouse

Based on the Homologene database [27], we identified orthology mappings between human and mouse genes, using the Homologene gene family IDs. Every individual human to mouse gene mapping was categorized as being part of a one-to-one, one-to-many, many-to-one, or many-to-many mapping between gene sets, based on the relative timing estimates for duplication and speciation events. Additionally, genes with no orthologs in the other species were categorized as one-to-none, many-to-none, none-to-one, or none-to-many mapping. We additionally curated the Homologene database to correct orthology relationships that were inconsistent with the literature, most notably the relationships in the MRGPR family. Ensembl Compara [109] gene trees were used to guide this process, and the final set of homology mappings used are presented beside abundance data for each gene in Sheet 1 of Supplementary Files 1 and 2. Correlational measures (Pearson Correlation Coefficient or PCC [81]) previously used in the literature [90] were calculated between the gene expression (uqTPM) vectors of human genes and their mouse orthologs across corresponding tissues, for those human and mouse genes that have a one-to-one orthology mapping and show either human or mouse DRG enrichment. We analyzed the empirical distribution of the DRG enrichment scores to identify hDRG enriched and mDRG enriched genes (DRG enrichment score > 0.29 in at least one species, corresponding to inclusion of all genes where uqTPM is highest in DRG across tissues) for all human and mouse genes, and restricted this gene set to genes with high correlation (determined from empirical distribution of the PCC) between human and mouse tissue expression panels.

### 2.4 Similarity analysis of gene and tissue expression profiles to identify gene expression signatures

The microarray datasets used were variations of the Affymetrix HG-U133A, and for each chip, the tool Oligo was used for microarray probe normalization [9], followed by quantile normalization across chips [8]. Averaging across replicates, a clear unimodal distribution for probe intensity was identified (Supplementary Fig. 3) for classifying probesets as expressed or unexpressed. The normalized intensity values are tabulated in Supplementary File 3. Since culturing cells can alter gene expression profiles [123], we ignore probesets that were not expressed in hDRG or TG and characterized which of the hDRG or TG expressed probesets were expressed in NHSCs or fibroblasts.

For RNA-seq, we analyzed only genes with polyA+ transcripts, and performing upper quartile normalization, we aimed to minimize the effect of technical variation in our data. Within human datasets, genes were analyzed for co-expression patterns across all analyzed tissues, to identify sets of genes with similar expression patterns and cluster them into “modules” [23]. We assigned each gene a digital “co-expression pattern”, based on whether the uqTPM in each tissue was >= 0.75 or not. The most frequent gene co-expression patterns were identified, along with TFs and other relevant genes in each module. The genes in the corresponding modules were then hierarchically clustered [23] and the resulting dendrogram optimally leaf-ordered on the genes [6] to depict the gene expression patterns across the set of analyzed human tissues. This allowed us to visualize relevant co-expression patterns and putatively identify regulatory circuitry underlying such co-expression.

We also performed hierarchical clustering of all human and mouse tissue transcriptome profiles, restricted to the set of genes to those with one-to-one orthology between human and mouse based on our analysis [27]. For the hDRG and mDRGs, all 3 individual replicates were individually used instead of a single tissue profile to identify the sample-to-sample variation relative to observed tissue-to-tissue variation. The set of genes used for this analysis was limited to genes with tissue-restricted expression in order to identify a clear tissue-specific gene signature [85].

### 2.5 Functional enrichment analysis

We identified human and mouse genes with relevant functional gene ontology descriptions, based on the Gene Ontology (GO) database [15]. Wherever possible, the set of human genes were augmented using the HGNC database [30]. The gene sets used are noted in Supplementary File 4. Functional enrichment analysis was then performed using Enrichr [11], allowing for multiple testing correction, with the Reactome database [17] being used for pathway analysis. In order to identify all known drug – gene product interactions for genes present in the conserved DRG enriched gene set, the Drug – Gene Interaction Database (DGIdb) [110] was used. Protein-protein interaction (PPI) were mined based on validated interactions of genes with relevant GO terms in the StringDB database [26]. The KEGG database and visualization tool [42] was used to analyze the neurotrophin signaling pathway, used with permission from Kanehisa Laboratories (Supplementary File 5).

### 2.6 *In silico* detection of mRNA axonal localization

We contrasted relative abundances of mRNAs in the hDRG from our own RNA-seq experiments to human tibial nerve (hTN) RNA-seq experiments from the GTex consortium [16]. Changes in gene expression between the hDRG and hTN can be brought about by region-specific gene expression in non-neuronal cells, subcellular localization in sensory neurons, variable composition of cell types between hDRG and hTN samples, differential expression between sensory (present in hDRG and hTN) and motor neurons (present only in hTN) or a combination of all of these. The hDRG: hTN fold change was calculated based on TPM values for all genes, and is presented in Sheet 1 of Supplementary File 2. We restricted our analysis to genes whose orthologs show differential expression amongst mDRG neuronal subpopulations, since such sensory neuronal subpopulation restricted expression typically involves neuron-specific expression [51].

## 3. Results

We introduce an open-data, searchable website for our analyses allowing users to visualize gene expression profiles (http://www.utdallas.edu/bbs/painneurosciencelab/DRGtranscriptome) across orthologous human and mouse genes. For reproducibility, code for performing calculation of the data underlying the figures: uqTPMs, and sample statistics for characterizing tissue specificity and conservation of expression are available in Supplementary File 6, and on our website.

### 3.1 Comparison of hDRG whole transcriptome profile with other transcriptome profiles

From a transcriptional point of view, the hDRG is a highly heterogeneous conglomerate of cells with multiple types of sensory neurons, Schwann cells, satellite glial cells, fibroblasts, and resident macrophages [72]. *In vivo* studies of heterogeneous tissues like the hDRG or hTN pose a special challenge, since mRNA species may be expressed specifically in individual cell types or more generically. We estimated the degree of overlap between the whole DRG transcriptomic profiles and some of its non-neuronal constituent cell types. We looked at assembled microarray datasets for human DRG, TG and cultured normal human Schwann cells (NHSC) and skin fibroblasts (FIBRO) (Fig. 2A). We identified 12,422 probesets that show expression in hDRGs. Of these, 11,407 (94.6%) are shared with the human TG, reiterating the overall similarity of the two tissues. Interestingly, we found that approximately 70% (8,666 / 12,422) of the DRG-expressed probesets show co-expression in either the NHSC or fibroblast dataset or both. The remaining 30% of the DRG-expressed probesets were putatively due to expression restricted to sensory neurons or SGCs and included nociceptor-enriched channels like SCN10A and SCN11A [107], and nociceptor-enriched transcription factors like PRDM12 [12] which are well characterized in mammalian DRGs. Our analysis indicates that cDNA libraries constructed from whole hDRGs have a strong non-neuronal component.

**Figure 2.**
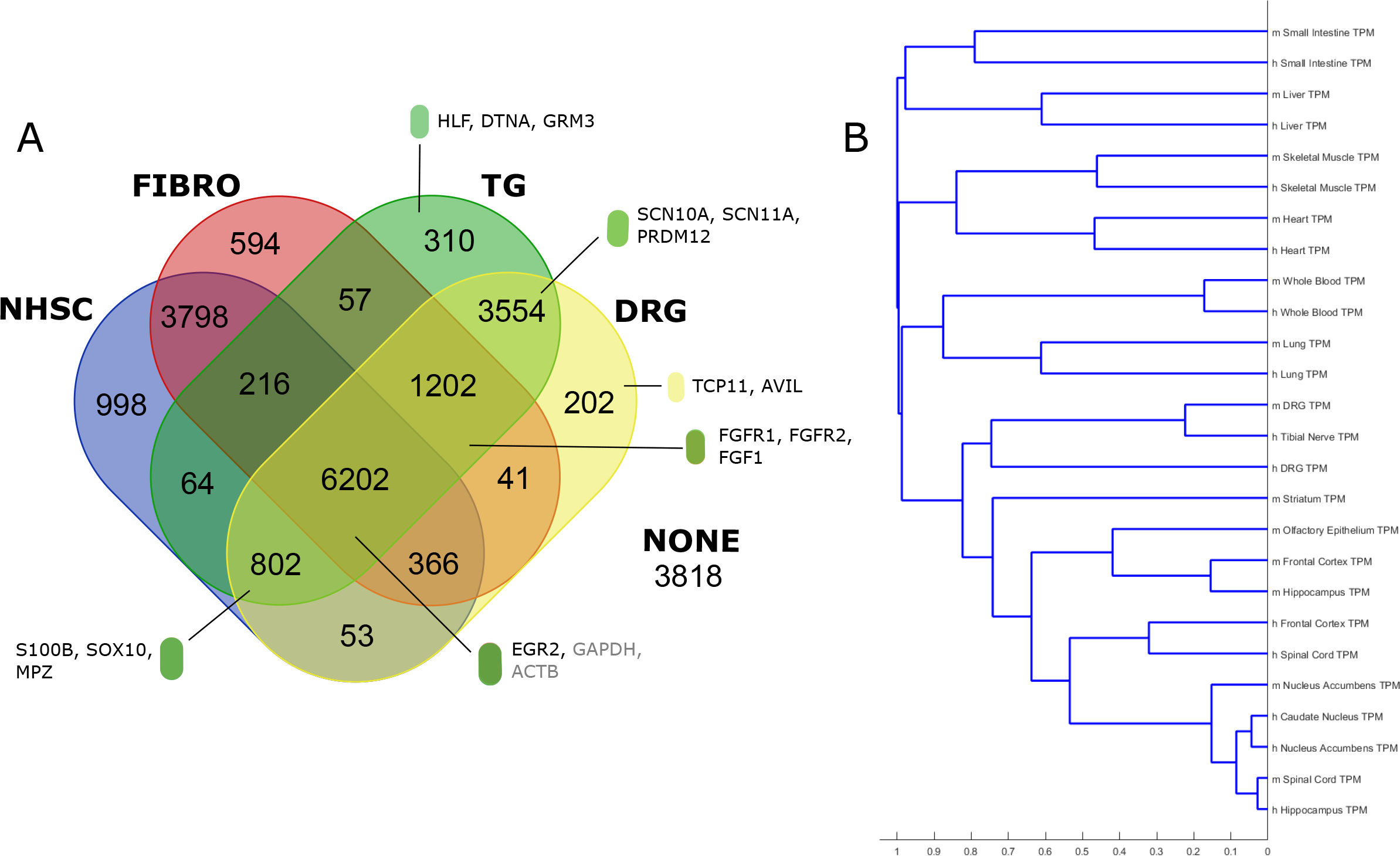
Cell-type specific and cell compartment specific gene expression in the hDRG. **(A)** Microarray analysis of human DRG, TG, cultured Schwann cells (NHSC) and fibroblasts (FIBRO) reveal that over 3500 probes are expressed *in vivo* in DRG and TG but not detected in Schwann cells or fibroblasts, suggesting regulatory programs driving gene expression specifically in sensory neurons and SGCs. **(B)** Clustering of human and mouse tissue transcriptomes based on a Pearson’s correlation coefficient (PCC)-based distance measure (1 – PCC) and average linkage for tissue-restricted genes.

To look at conservation of gene expression across tissues in humans and mice in a transcriptome-wide fashion, we performed hierarchical clustering of the RNA-seq transcriptomes (Fig. 2B). We find that for non-neural tissues, homologous human and mouse tissues cluster in pairs. The hDRG replicates, mDRG replicates and hTN sample also cluster together, showing a distinct PNS gene expression profile, and low sample-to-sample variation in our hDRG and mDRG samples. Our findings agree with meta-analyses that suggest distinct signatures for human sensory tissues and brain tissues [100], and that inter-study distances among homologous tissue transcriptomes are typically less than intra-study distances between different tissues [99]. However, inside the CNS tissue cluster, we find that the human samples and mouse samples do not cluster according to brain regions, and we attribute this to well characterized evolutionary divergence of the CNS transcriptome in humans with respect to other primates and rodents [61; 76] rather than batch or laboratory effects as the subclusters are not determined by source laboratory of the samples. Preclinical rodent models have been shown to mimic human response in several domains [100]. Our analysis identifies a distinct, evolutionarily conserved tissue-specific signature for healthy human and mouse PNS tissues, opening the door to the follow up question of how faithfully rodent pain models correlate with human pain signatures at the molecular level.

### 3.2 Co-expression patterns and regulatory programs shaping the hDRG

Based on our binary co-expression pattern for each coding or non-coding human gene, we found that 22,140 genes were expressed in one or more tissues, with 12,462 genes being generically expressed across the reference tissue panel based on their entropy (Fig.s 3A and 3B). Of the remaining 9,498 genes, we found a power law-like distribution for frequencies of possible co-expression patterns across the tissues we queried (Fig. 3C), suggesting that a few dominant regulatory paradigms primarily shape the hDRG transcriptome. Out of the top 25 most frequent patterns (5,542 out of the 9,498 tissue-restricted genes in our analysis), 15 correspond to enrichment or de-enrichment in subsets of non-neural tissues, 6 to enrichment in subsets of CNS tissue (Fig. 3D), and the remaining 4 affecting hDRG expression. The most common co-expression pattern of these was the set expressed pan-neuronally (Fig. 3E), the basis of a neuronal gene signature [24]. Previous analyses with hDRG microarray datasets were limited to analyzing a set of predefined probesets, and were unable to identify a broad pan-neuronal transcriptome signature including the hDRG [88; 98]. In our analysis, pan-neuronal genes include splicing factor RBFOX3, ion channels (SCN1A, KCNK1), tyrosine receptor kinase NTRK2, cell adhesion molecule (CAM) gene DOCK3 and the early neural lineage TF SOX1 [82]. The second relevant gene co-expression module consists of genes which are downregulated in the hDRG and spinal cord but expressed in the rest of the CNS (Fig. 3F), potentially driven by TFs like HOXA3, HOXA4 and HOXB3 involved in rostral-caudal patterning [73] and are common to these two adult human tissues but not more rostral parts of the nervous system. The TF MYT1L is downregulated in hDRG and spinal cord with respect to the rest of the CNS. A third prominent co-expression pattern identifies genes enriched primarily in the CNS but undetectable in the hDRG (Fig. 3G). These include CNS-expressed genes like SLC32A1 [75] and GLRA2 [122], known to be suppressed in the DRG, the metabotropic glutamate receptor GRM5 (linked to postsynaptic plasticity in the spinal dorsal horn [36]), RNA binding protein PABPC1L2A, serotonin 2C receptor HTR2C, and TFs OLIG2 (present in astrocyte and oligodendrocyte cell types, and subpopulations of motor neuron [56]) and POU3F4. The fourth prominent co-expression pattern was primarily enriched in the hDRG (Fig. 3H), with the gene set explored in more detail in Section 3.3.

**Figure 3.**
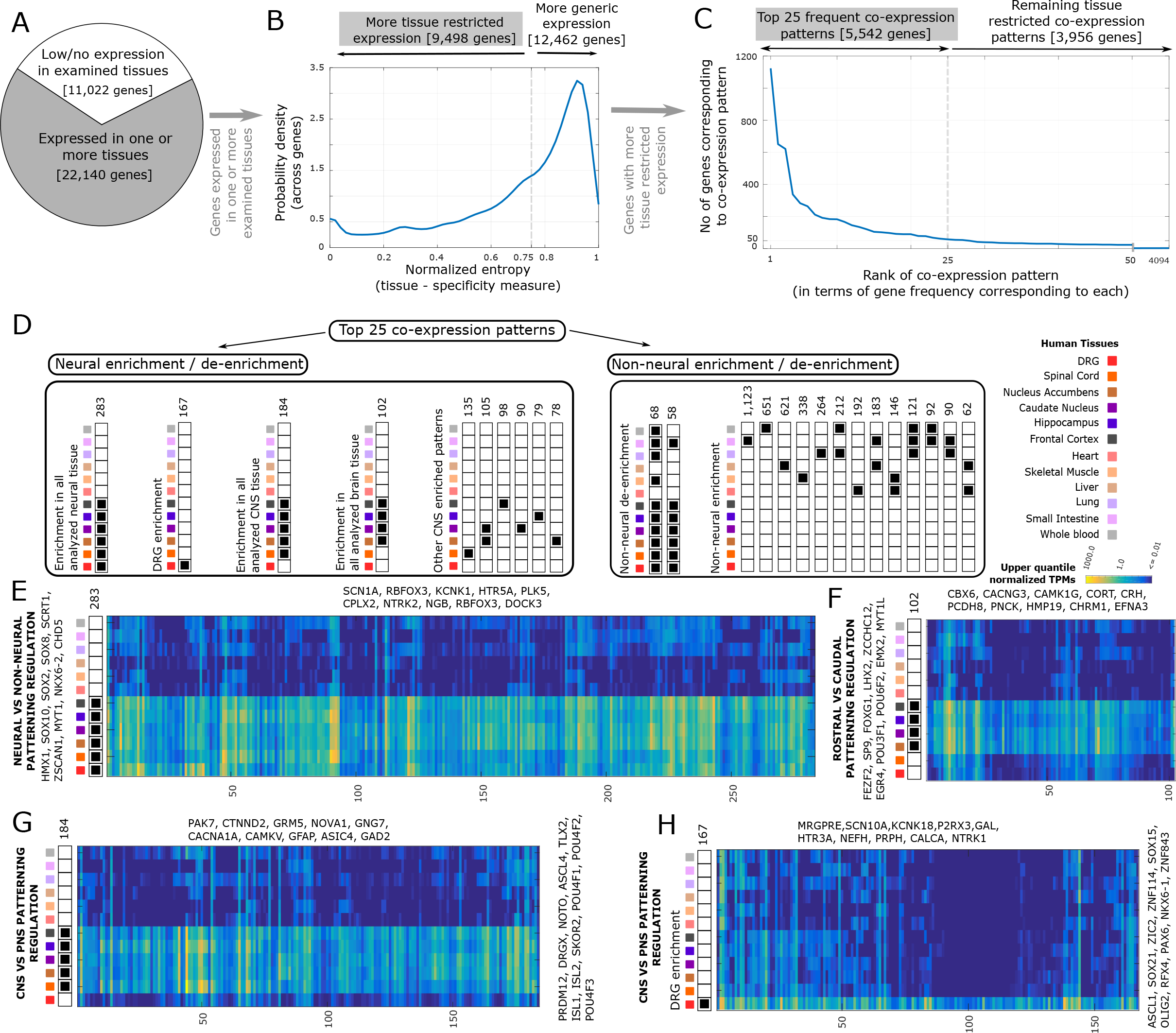
Identification of common tissue-restricted human gene co-expression patterns in our analyzed tissue panel. **(A)** ~22,000 out of ~33,000 genes are expressed in at least one tissue based on our digital co-expression pattern **(B)** 9,498 genes showed tissue-restricted expression (based on a normalized entropy measure of tissue diversity) **(C)** 25 co-expression patterns are sufficient to account for 5,542 of these genes, in a power law - like distribution. **(D)** 25 common co-expression patterns. Heatmaps with hallmark genes and potential transcriptional regulators which are differentially expressed in **(E)** neural versus non-neural tissue **(F)** brain tissues versus remaining tissue **(G)** CNS versus non-CNS tissue **(H)** DRG versus remaining tissues.

To identify a more comprehensive picture of the regulatory forces driving gene expression in the hDRG, we catalogued all hDRG-enriched and neurally enriched TFs (heatmaps for human and orthologous mouse gene expression in Fig. 4A). Multiple evolutionary developmental studies have been performed [12; 53; 69] suggesting an important role of TLX3, RUNX1, DRGX, POU4F1 and PRDM12 in mouse, and *Xenopus* sensory tissue development. We find that most of the studied regulatory TFs have conserved DRG-enriched gene expression in adult humans and mouse, suggesting that regulatory interactions are conserved through evolutionary history. The most well-studied among these TFs is possibly PRDM12, which is essential for mammalian DRG development and pain sensation [12]. The ancient origins of somatic and visceral neurons is well documented in bilaterans (arthropods and vertebrates) [74], and among the transcriptional determinants of such neurons are DRG-enriched TFs DRGX, POU4F1 and POU4F2 (both of the BRN family). Based on mDRG scRNA-seq data [106], several of the DRG-enriched TFs (HOXD1, PRDM12, POU4F2, POU4F3, ISL2) are restricted to subpopulations of adult mDRG neurons while POU4F1 and ISL1 are more widely expressed across subpopulations. The TFs may thus collaborate in subpopulation-specific ways to drive regulatory programs specific to each of the neuronal subpopulations. *In vitro* differentiation of human pluripotent cells using POU4F1 (and either NGN1 or NGN2) [7], reprogramming of human fibroblasts using a TF cocktail of multiple subpopulation-restricted TFs including ISL2 [111], and characterization of sensory neurons induced from human Pluripotent Stem Cells uniformly expressing ISL1 and POU4F1 [10] suggests that a transcriptional program similar to mDRG neurons may be at work in humans.

**Figure 4.**
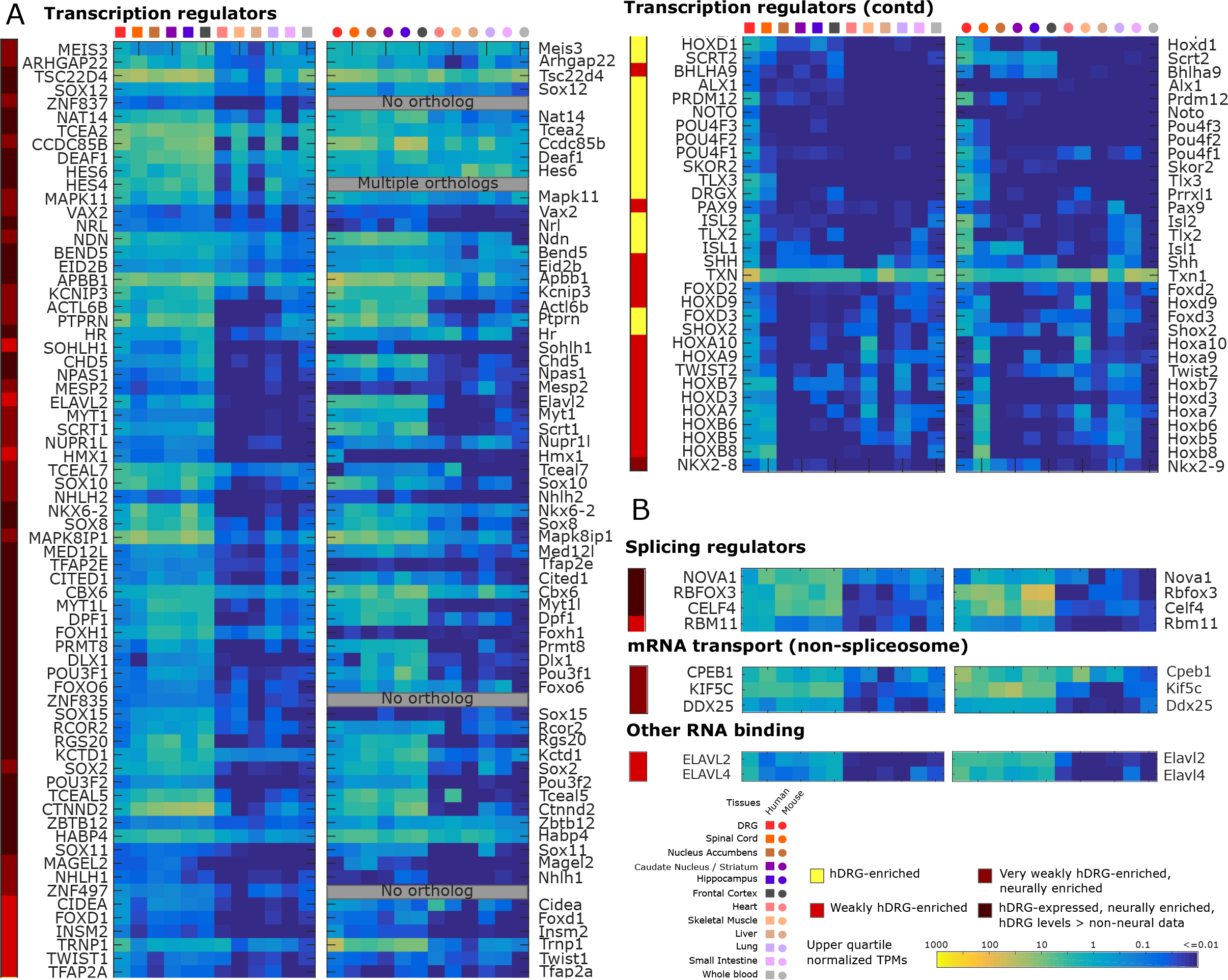
Identification of hDRG enriched genes. **(A)** Flowchart for identifying DRG-enriched genes with conserved expression patterns in human and mouse. **(B)** Estimated density function in DRG-enriched genes for Pearson’s correlation across vectors of relative abundance in orthologous tissues. **(C)** The set of hDRG and mDRG enriched genes (based on the DRG enrichment score) with one-to-one orthology and correlated gene expression abundances (based on Pearson correlation) across our panel of analyzed tissues. Many of the 81 genes belong to gene families known to play important functional roles in nociceptors, including GPCRs, ion channels, TFs, and transmembrane proteins. **(D)** Assessment of expression of hDRG genes in mDRG scRNA-seq data [107] for prediction of potential sensory neuronal subtype expression.

Several hDRG-expressed TFs, co-factors, splicing regulators and mRNA binding proteins (both spliceosomal and non-spliceosomal) show neural enrichment, but are only weakly hDRG-enriched (Fig.s 4A and 4B). Several of these are potentially involved in hDRG function, and are elucidated in Table 2.

These analyses give a global view of DRG gene expression in humans compared to a number of other tissues, highlighting that the DRG expression profile is distinct from CNS gene expression patterns [88; 98] driven apart during development by well-characterized CNS-specific TFs like POU3F3, and SOX21 and DRG-specific TFs like PRDM12, DRGX and TLX3. While DRG-specific [53; 69; 86] and CNS-specific TFs [65; 112] have well-appreciated roles in development, their persistent expression into late adulthood in humans has previously not been appreciated, suggesting a role of cell-type maintenance alongside that of the well-characterized developmental lineage drivers. Additionally, multiple DRG-enriched TFs (ISL1, ISL2, DRGX) play a dual role in developmental axonogenesis and adult axon regeneration. Human DRG-specific TFs are also likely involved in regulating the transcription of hDRG-specific genes, although specific TF targeting of DRG-specific genes underlying the regulatory program are not well understood except for a small subset of genes profiled in evolutionary development studies [12; 53; 69].

### 3.3 hDRG and mDRG enriched genes and a conserved evolutionary signature

Based on the DRG enrichment score, we identified 140 protein coding genes to be hDRG-enriched, and 141 protein-coding genes to be mDRG-enriched. Of these 128 hDRG-enriched and 119 mDRG-enriched genes (identified based on Section 2.2, Fig. 5A) have one-to-one orthologs in the other species. We then looked at the correlation between the human and mouse ortholog uqTPM expression across the analyzed tissue panel to identify if the gene in the other species also had a DRG-enriched expression profile, reported in Table 3. When we analyzed the distribution of correlation values, we found a large population of highly correlated relative abundance, suggesting that a large fraction of these genes are under negative evolutionary selection in their regulatory regions, causing conserved gene expression profiles across tissues (Fig. 5B). 81 out of 128 (63.3%) human genes have correlation scores above 0.68 (the peak in the distribution before the primary mode). All 81 of these genes in both species have the highest relative abundance in DRG across the panel of analyzed tissues, showing a clear conservation of DRG-enrichment. Among the genes we identified with a common pattern of DRG enrichment in mouse and human (Fig. 5C), are several that have not been previously studied in DRG biology. These include UCHL1 (neuroepidermal marker of itch [49]), SKOR2 (a developmental co-repressor in cerebellum [70]), TRIM67 (shown to be involved in RAS-mediated signaling in the cerebellum [118]), BET3L (particle transport complex member identified in mDRG single cell studies [51] but not functionally studied), and TLX2 (whose paralog TLX3 is well characterized as a key DRG-enriched TF).

**Figure 5.**
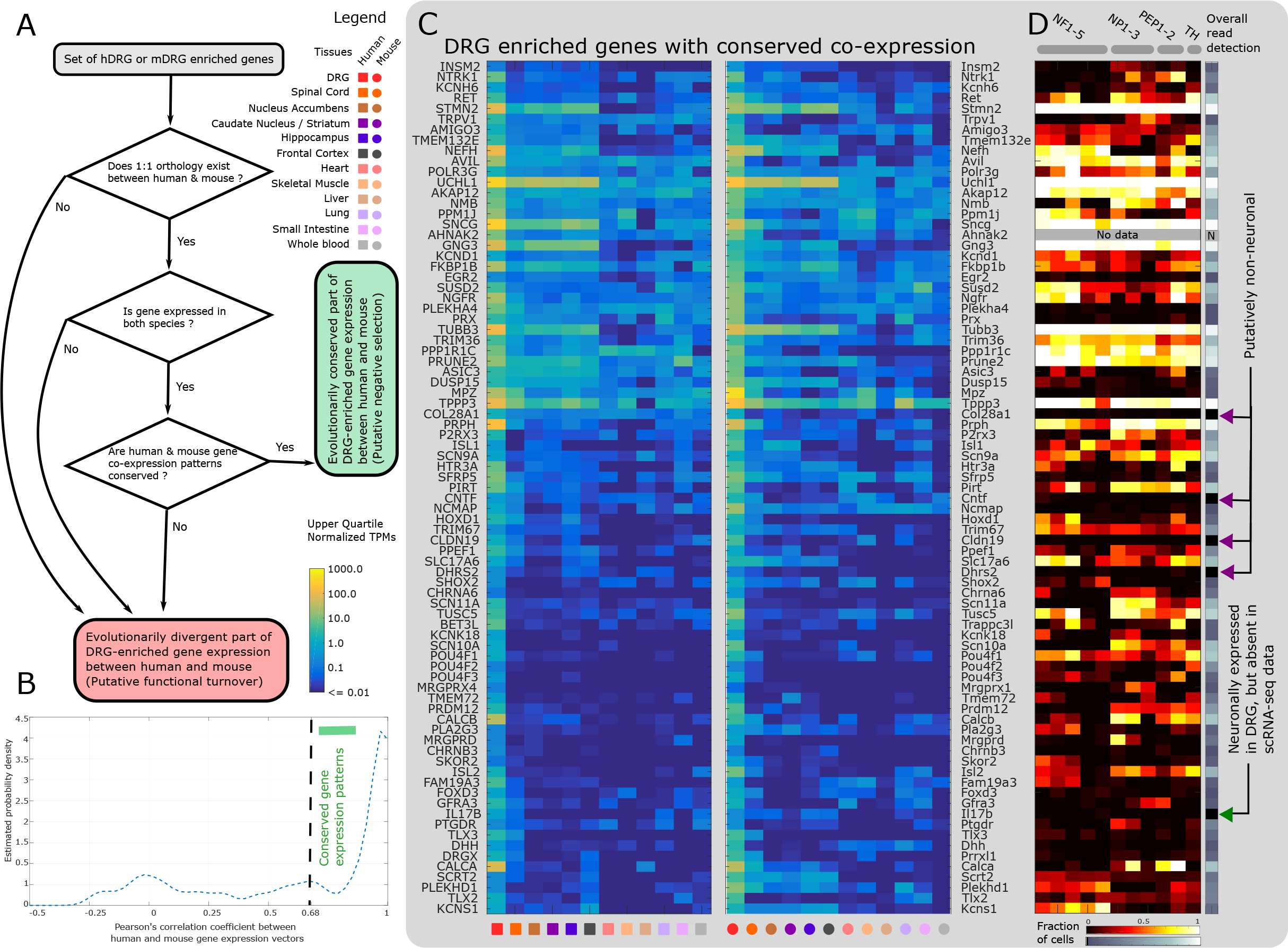
hDRG-enriched genes that potentially regulate the transcriptomic landscape. Orthologous human and mouse gene expression in the corresponding tissues is depicted as heatmaps. Differentially expressed TFs **(A)**, splicing factors, mRNA transport molecules, and RNA binding proteins are shown **(B)**.

Several DRG-enriched genes show subpopulation restricted expression in the mDRG (expression profiles in mDRG neuronal subpopulations based on Usoskin *et al* [106] are shown for all DRG-enriched genes in Fig. 5D and Table 4), raising the possibility of a similar expression profile in sensory neuronal subpopulations in the hDRG. Among the more well-studied genes are NTRK1, TRPV1, and CALCA (mRNA found in both peptidergic and non-peptidergic neurons), ASIC3 (neurofilament and peptidergic neurons), MRGPRD (non-peptidergic neurons), and PRDM12 (peptidergic, non-peptidergic and TH-positive neurons).

We also identified a gene set which have undetectable gene expression in mDRG scRNA dataset, many of which are known to be expressed in mammalian Schwann cells, detailed in Table 5. Known peripheral SGC markers are either weakly expressed in our hDRG datasets (like PAX3 or PAX7) or also expressed in other tissues (like GFAP) and are thus not DRG-enriched. Fibroblast markers like S100A4 are highly expressed in DRG, but are also generically expressed, leading to low DRG-enrichment scores.

We identified that many of the gene products of hDRG-enriched genes can be assigned a handful of molecular roles including transcriptional activators or repressors, receptor and receptor binding, kinases or phosphatases, ion channels, and transmembrane domain containing proteins. Based on the Enrichr tool [48], we performed enrichment analysis for biological processes in the Gene Ontology (GO) database [3] (Table 6) and molecular pathways in the Reactome database [17] (Table 7) that were enriched in our gene set (Fisher’s exact test, *p* < 0.01 after multiple testing correction). Gene or protein functional ontologies are well known to be incomplete [40], necessitating us to look at DRG-enriched genes in non-enriched ontology terms and the literature, and showing the diverse roles played by these gene products. Several of these are already well studied in DRG biology. CALCB is a paralog of CALCA with similar function. MRGPR family members are known to play a key role in itch sensation [52]. PIRT is a well-known regulatory subunit of TRPV1 [43], while PRDM12 is a key TF in neural sensory lineages [12]. Querying the Reactome database (Table 7) yielded enrichment for acetylcholine and nicotinic receptor signaling centered around 3 specific receptor subunits, β3, α6 and α9. The CHRNA6 gene was first identified as a gene potentially involved in mechanical allodynia in mouse, leading to an identification of a novel interaction between α6 nicotinic receptors and P2X3 ion channels that leads to inhibition of P2X3 channel activity [113]. CHRNA9 (encoding the α9 nicotinic receptor subunit) was recently implicated in development of chemotherapy induced peripheral neuropathic pain using an antagonist of α9 and α10 nicotinic receptors [80]. While more work is needed to validate α9 as the mechanistic target for this compound, the very strong enrichment of CHRNA9 in human DRG makes it an interesting target for this prominent form of neuropathic pain.

More interestingly, we identified many genes whose functional roles in DRG have not been well studied (Fig. 5C). PPP1R1C, PPM1J, and PPEF1 are phosphatases: PPM1J is known to regulate neurite growth [2], PPEF1 interacts with Calmodulin in a Ca^+2^ dependent manner [50], and DUSP15 regulates the JNK pathway [93]. AMIGO3 [47] is involved in axonal tract development. SUSD2 is known to regulate neurite growth in hippocampal cultures [68]. TPPP3, PLA2G3, and TRIM36 [97] (along with TUBB3) are associated with the microtubule cytoskeleton. AKAP12 is a kinase scaffold involved in regulating cAMP biosynthesis and signaling. TUSC5 is expressed in the PNS and adipose tissue, and potentially performs shared adipose-nervous system functions [77]. PTGDR and IL17B are known to be involved in inflammation response, and POLR3G is also involved in cytokine production and immune response. BET3L is part of the TRAPP protein trafficking complex, while NMB is a neuropeptide involved in nociceptive signaling [64]. Additionally, we have a very limited understanding of the role of some DRG-enriched neuronally expressed genes. INSM2, SHOX2, SCRT2, and SKOR2 are development-related transcriptional regulators but their regulatory function in adult DRG is not known. The neuronal roles of PLEKHD1, AHNAK2, PRUNE2, FAM19A3, and the transmembrane proteins TMEM72 and TMEM132E are also unclear. These genes create a rich set of novel targets for potential exploration in the pain neurobiology space.

Two previous studies have characterized DRG-enriched genes in humans, but were limited in their scope. Flegel *et al* [25] analyzed hDRG-enriched genes by chronicling known functionally important gene families in the DRG, and Sapio *et al* [91] analyzed hDRG-enriched genes from the perspective of sensory neuropathies. In contrast to these, we perform an unbiased, multifaceted enrichment analysis to characterize this gene set. While some of these genes were previously known to be DRG-specific (like TRPV1, MRGPRD, and SCN10A), we identified several genes (like QRFPR) that have not been previously shown to be part of a DRG-enriched transcription regulatory program.

### 3.4 Identifying potential pharmacological targets among tissue-restricted genes in the hDRG

Since a primary goal of our analysis is the identification of potential therapeutics, we performed additional analysis on our DRG-enriched and neurally enriched gene sets to mine for known or new drug targets. We first used our hDRG-enriched gene set to explore the drug – gene product interaction database DGIdb [110] for targets of known drugs (Table 8). This identifies genes for which ligand interactions are known, potentially identifying new drugs that can be repurposed or redesigned to target pain. We identified a subset of gene-ligand interactions from DGIdb but these were mostly well-known pain targets with ligands moving into or already in clinical trials. For instance, CALCA (*CGRP*), P2RX3 (*P*_*2*_*X*_*3*_), NTRK1 (*TrkA*), SCN10A (*Nav1.8*) and SCN11A (*Nav1.9*) are all well-known genes involved in pain, most of which have robust ongoing clinical candidate development. Others in this list included nicotinic receptors that were mentioned above but most of the ligands identified lack specificity for these targets. The HTR3A gene, which encodes the 5-HT3 ligand gated ion channel, is enriched in hDRG and many ligands of this receptor subtype are known. Despite a strong case for the role of this receptor in chronic pain states [44], the pain-relieving potential of antagonists of this receptor have never been clarified in humans, although one small clinical trial in neuropathic pain suggests efficacy of antagonists [57]. Only one gene in this list clearly stood out as novel, QRFPR, which encodes the pyroglutamylated RFamide peptide receptor. This receptor has a known specific ligand but its role in sensory biology and/or pain has not been assessed.

The short list of known ligand to gene interactions identified from the DRG-enriched dataset suggests that many DRG-specific genes have not been explored as drug targets. We thus characterized several pharmacologically important gene families including kinases, phosphatases, ion channels, GPCRs, neuropeptides and associated receptors, and cell adhesion molecules. We also considered that drug targets can include drugs that do not cross the BBB and we therefore included hDRG-expressed genes that had strong neuronal expression but did not show expression in non-neuronal tissues. These genes may be targets for drugs that do not cross the BBB, if DRG targeting is the desired outcome.

Among ion channels, we identified several ion channels with well-known DRG-specific expression (SCN10A, TPRV1, ASIC3 and P2X3) but also identified several others (Fig. 6A), such as nicotinic receptor subunits α6 and α9 (mentioned above) and potassium channels (KCNK18, KCNS1 and KCNH6). A much larger set of ion channel genes showed enrichment in neuronal tissues with preserved expression in hDRG, and in many cases also in mDRG. Among these were many additional potassium channel genes and strong expression for kainate receptor subtypes, in particular in the hDRG. Several neuropeptide and neuropeptide receptors have been targets of new drugs developed in the past decade [31]. Some candidate drugs targeting these molecules have not been studied due to low bioavailability (because of BBB and other factors) [22], which is not necessarily a challenge when targeting the DRG. NMB, CALCA, CALCB, and GAL were identified to be hDRG-enriched, as was the receptor QRFPR, which was mentioned above. GLRA4, which is a glycine receptor subunit, was expressed in hDRG but not mDRG (Fig. 6B). Cell Adhesion Molecules (CAMs) are also increasingly viewed as pharmacological targets due to their involvement in processes like inflammation, and cell signaling [45]. We identified several neuronally enriched CAMs like CNTN1, NLGN3, NRCAM, and LRRC4C, but none of these were specific to hDRG (Fig. 6C). DRG-enriched MPZ and CLDN19 are expressed in Schwann cells, but have gene products that are present at neuron - Schwann cell junctions.

**Figure 6.**
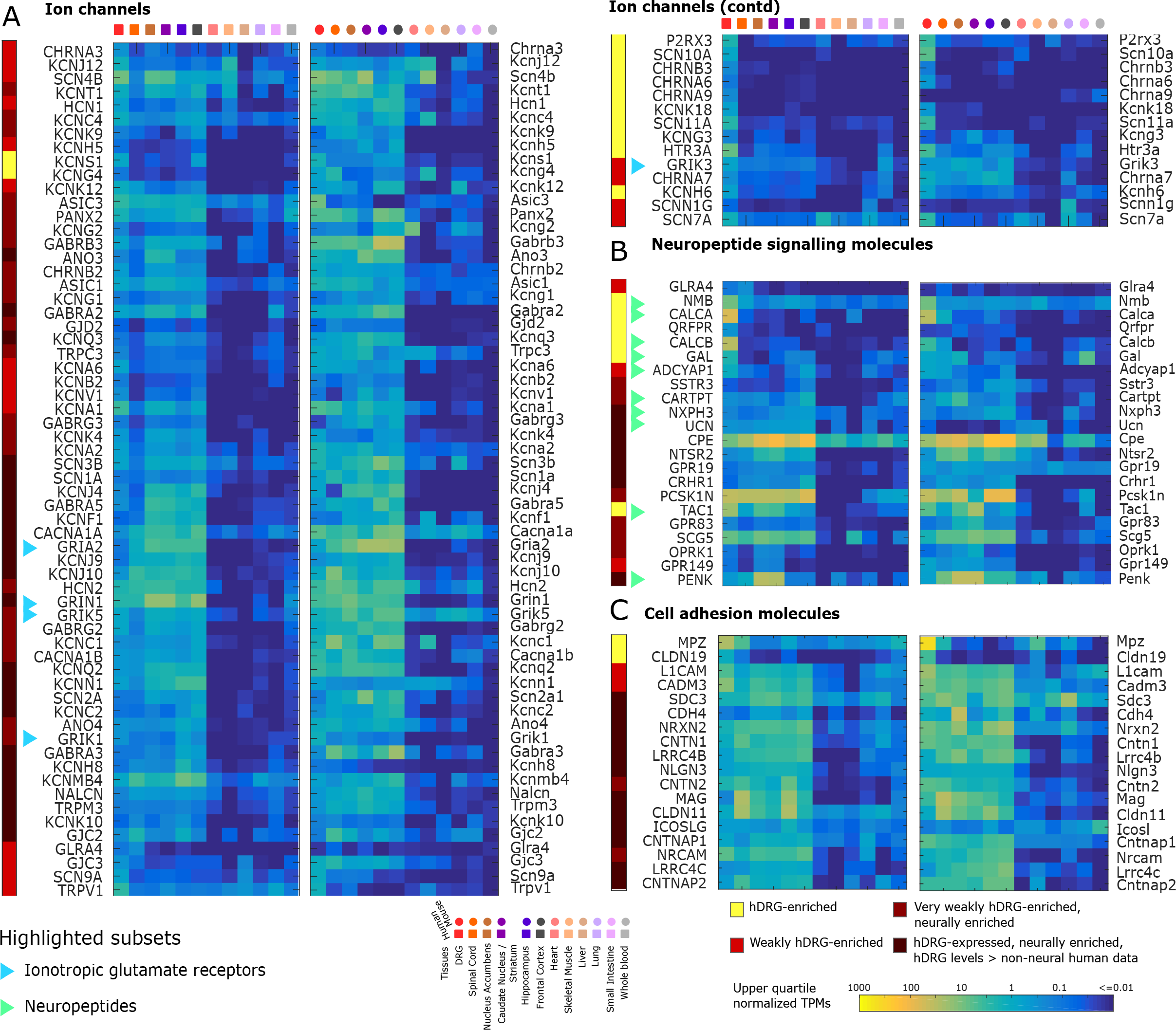
hDRG-enriched genes for pharmacological targets: ion channels, neuropeptides and cell adhesion molecules. Orthologous human and mouse gene expression in the corresponding tissues is depicted as heatmaps. Differentially expressed ion channels **(A)**, neuropeptide signaling **(B)** and cell adhesion molecules **(C)** show several candidate genes for pharmacological targeting, including several neuropeptides.

Among GPCRs, the Mas-related GPCR family (including known DRG specific genes in mouse and human) showed DRG enrichment (as expected) as did several other GCPRs including QRFPR, CCKAR and F2RL2 (Fig. 7A). We also identified several DRG-enriched (OR7E101P, OR51E2) or pan-neurally enriched (OR2L13, OR7A5) olfactory receptors. While we cannot assess if they are enriched in human olfactory epithelium due to a lack transcriptome data for this tissue, we find that the mouse orthologs for OR7A5 and OR51E2 are not expressed in the mouse olfactory epithelium (Sheet 1 of Supplementary File 2). While the role of olfactory receptors in non-olfactory tissue is under debate, some have been shown to be upregulated in nerve injury models [1], suggesting that these receptors may have functions outside of their canonical tissue. A broader list of GPCRs were expressed in hDRG and also found in other CNS tissues. Interestingly, these included a large number of orphan GPCRs, suggesting an unmined pharmacological resource for sensory and pain research. Finally, we assessed kinases (Fig. 7B) and phosphatases (Fig.7C). Notable among the kinases were NTRK1 and RET, receptor tyrosine kinases for NGF and GDNF, respectively, both of which are well-known to be DRG-enriched in adult human and mouse. Another weakly enriched receptor tyrosine kinase in hDRG was INSRR, an insulin receptor related receptor. All other kinases we identified were expressed in hDRG but also in other human CNS regions. Several neuron-enriched phosphatases were also detected, including tyrosine phosphatases PTPRN, PTPRN2, and PTPRZ1 but none of these showed a clear enrichment in hDRG.

**Figure 7.**
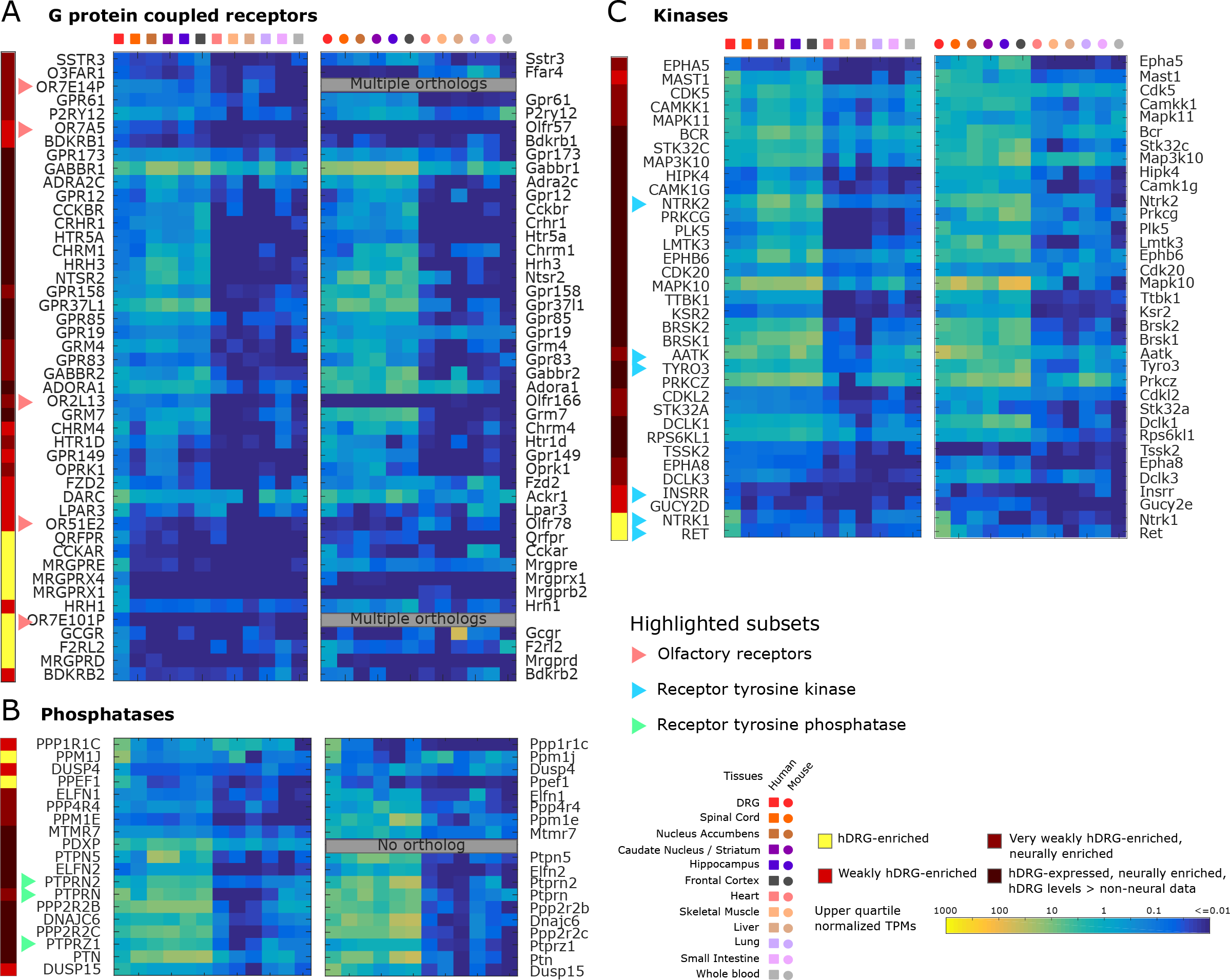
hDRG-enriched genes for pharmacological targets: GPCRs, phosphatases and kinases. Orthologous human and mouse gene expression in the corresponding tissues is depicted as heatmaps. Differentially expressed GPCRs **(A)**, phosphatases **(B)** and kinases **(C)** show several candidate genes, including several receptor tyrosine kinases and phosphatases, and olfactory receptors for pharmacological targeting.

To delve deeper into identifying DRG enriched genes as putative therapeutic targets, we looked at experimentally validated protein - protein interactions (PPI) in StringDB [26]. We limited this search to interactions within and between the set of DRG-enriched genes and the gene families with drug targeting potential, in a bid to identify pathways with strong therapeutic potential. Based on this network (Supplementary Fig. 4) and intersectional analysis with the KEGG pathway database [42], we identify several well understood pathways whose components are partially identified, including the neurotrophin signaling, cAMP signaling, retrograde endocannabinoid signaling, and axon guidance pathways. We detailed the neurotrophin signaling pathway (Fig. 8A), with focus on signaling through the NGF receptor since NGF-targeted therapies are in clinical trials for pain but have an uncertain future due to rare, severe side effects [5]. We find that most gene level abundances for the corresponding proteins in this signaling pathway are present in the hDRG and some are either hDRG-enriched (NTRK1, NGFR), or weakly hDRG-enriched (MAPK11, SHC2, ARHGDIA, ARHGDIG, Fig. 8B). These NGF signaling pathway components may be drug targets to manipulate this pathway in the hDRG with relative specificity.

**Figure 8.**
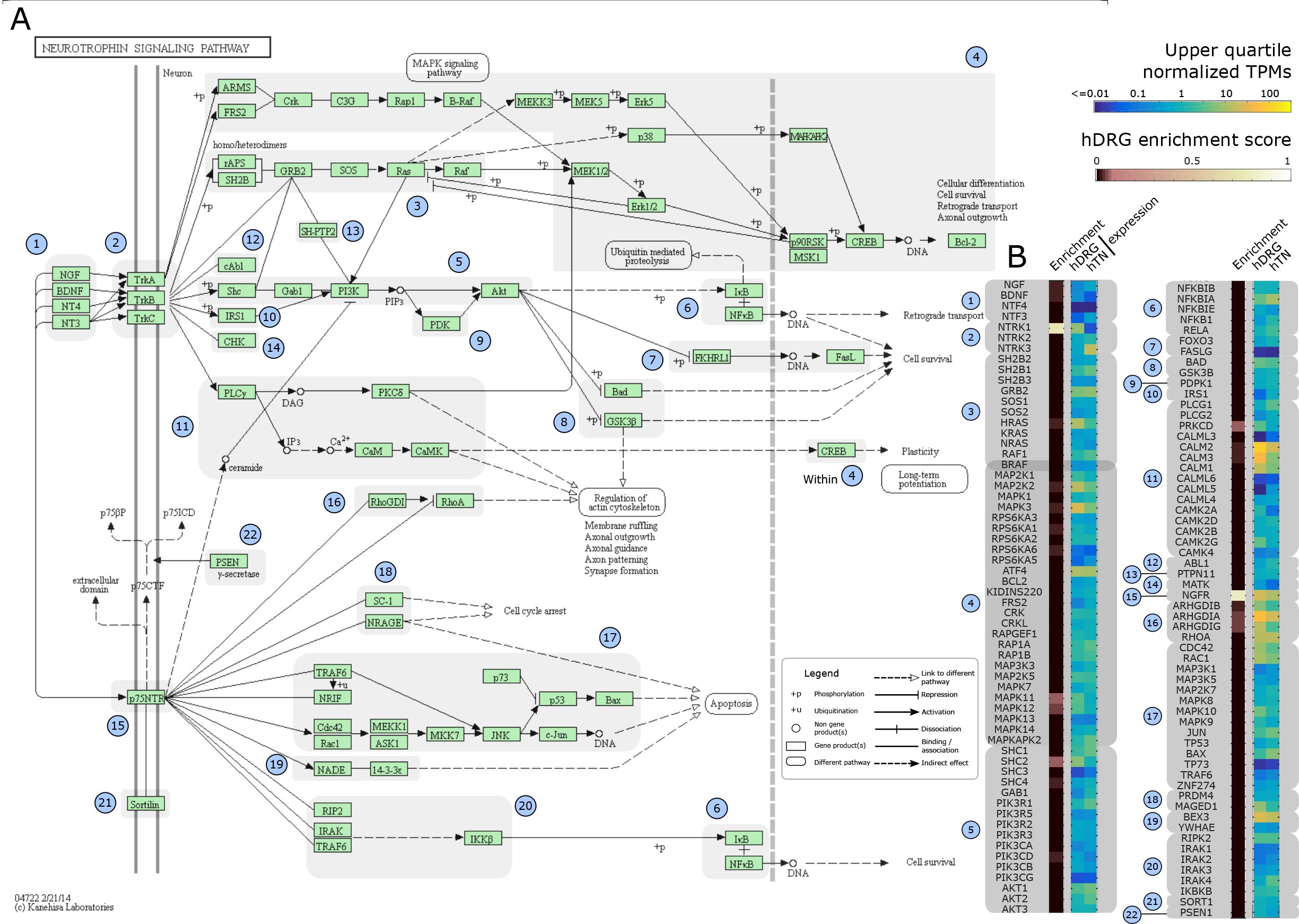
Enrichment of NGF/trkA signaling components in hDRG. **(A)** The NGF neurotrophin signaling pathway, based on the KEGG database, with permission from Kanehisa Laboratories (who retain copyright), **(B)** Heatmaps showing hDRG and human tibial nerve (hTN) expression and hDRG-enrichment for members of the pathway. Several signaling molecules in this pathway are enriched in the hDRG compared to other tissues analyzed in this study.

### 3.5 Putative axonal localization of mRNAs in hDRG sensory neurons

MRNA may be specifically compartmentalized inside somatic or axonal compartments of very long pseudounipolar sensory neurons. Gene expression changes in proximal versus distal portions of peripheral nerves in nerve-injured rats [39] have shown a clear signal of differential gene expression, and abundant evidence links axonally localized mRNAs in mouse DRG neurons to axon growth, regeneration and nociceptive plasticity [41; 59; 60; 63; 114]. Differential gene expression between peripheral nerves and the DRG in humans have only been characterized with respect to peripheral neuropathies [91], and putative localization of mRNAs to the axons of hDRG neurons has not been examined in general. We contrasted relative abundances of mRNAs in the DRG from our own RNA-seq experiments to human tibial nerve RNA-seq experiments from the GTex consortium [16]. Strikingly, we find fewer than 2,000 genes were detectable in one tissue and not the other, suggesting widespread axonal mRNA transportation (Fig. 9A).

**Figure 9.**
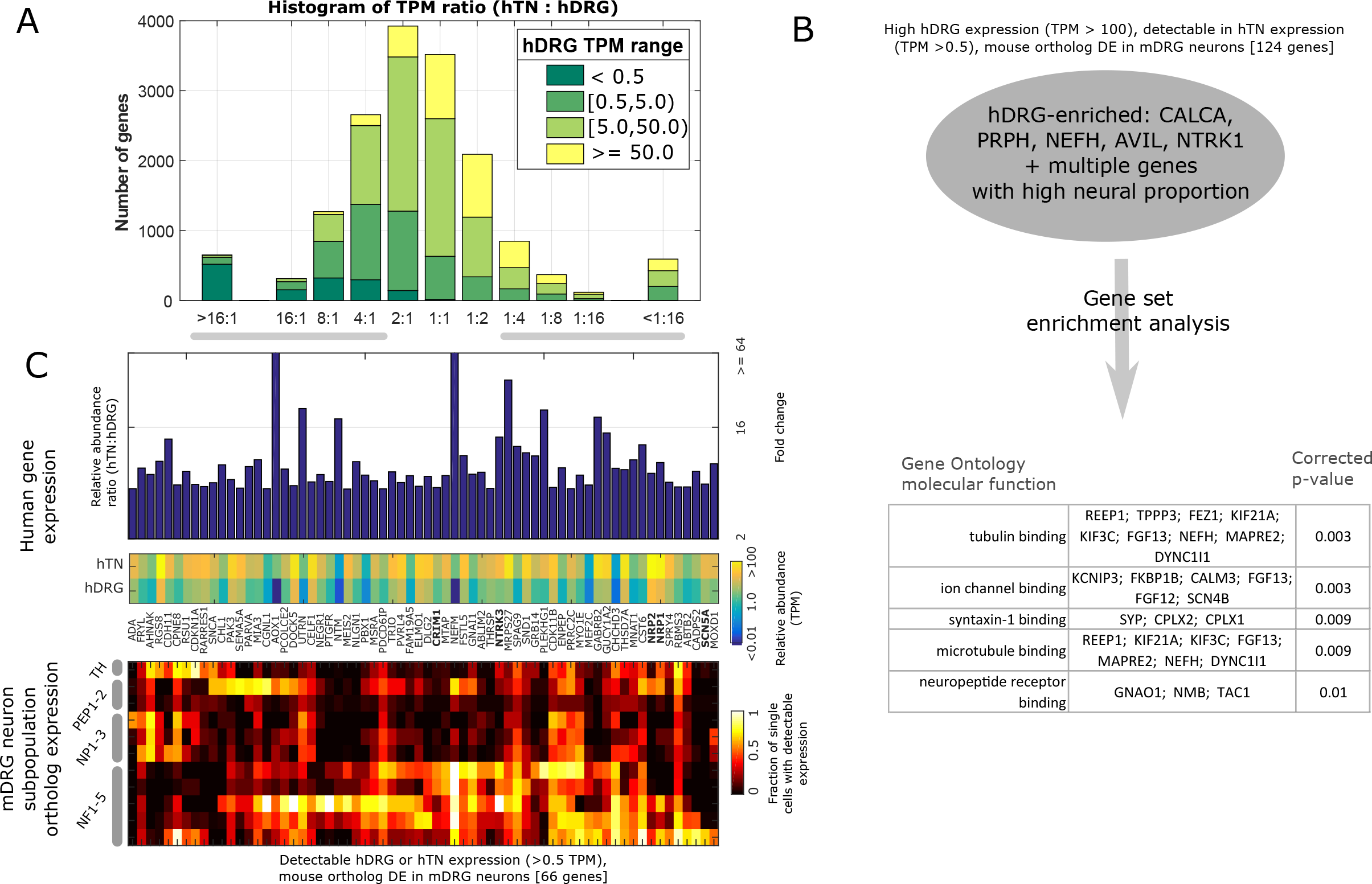
Comparison of gene abundances in hTNs and hDRGs identifies potential pervasive transport of hDRG mRNAs. **(A)** Distribution of hTN: hDRG relative abundances showing high overlap of transcripts **(B)** GO term enrichment for genes enriched in hDRG with respect to hTN transcriptome, restricted to genes whose orthologs were detected in specific subpopulations of mDRG **(C)** mDRG neuronal subpopulation restricted genes whose human orthologs are enriched in hTN versus hDRG.

We found that membrane protein PIRT (regulatory subunit of TRPV1) [43; 101] is localized only to the hDRG with no detectable expression in the hTN. Additionally, we noticed several sensory neuron-specific genes like MAS-related GPCRs MRGPRX1, MRGPRX3, and MRGPRE have downregulated or undetectable gene expression levels in hTN, but are detected at high levels in DRG, suggesting that these mRNAs are not axonally transported. These make them good candidates for neuronal subpopulation level cell body biomarkers for mRNA *in situ* hybridization studies.

The capacity for injury-induced axonal regeneration and pathfinding is a hallmark of adult PNS neurons [120]. We find several tissue repair associated genes to be more abundant in the hTN with respect to hDRG. The mouse RhoA effector Rock has been shown to be important in regulating axonal pathfinding [115] and to be an axonally targeted mRNA. In hDRGs, ROCK1 and ROCK2 expression were upregulated in hTN suggesting anterograde axonal targeting of mRNA for efficient local translation. Axon guidance receptors NRP1 and NRP2 were similarly more abundant in hTNs with respect to hDRG. Transcription factor protein sequences typically contain NLS sequences that promote retrograde transport to the nucleus. It has thus been hypothesized that anterograde transport of mRNA to axons, stimulus-induced local translation, and retrograde transport of protein to the nucleus is a potential mechanism for growth cone, or nerve ending to nucleus signaling [38] and similar signaling may promote nociceptor sensitization [60]. Several TFs that are known to be involved in axonal regeneration, including TP53, ATF3, FOS, JUN and STAT3 [67] all have higher relative abundance in the hTN over hDRG.

In order to broadly look at genes that were primarily expressed in sensory neurons in the hDRG and hTN, we further limited our analysis to orthologs of genes that are differentially expressed in mDRG neuronal subpopulations (Fig. 9B and 9C). Among genes detectable in the hTN, but more abundant in the hDRG, we performed functional enrichment, and identified gene sets corresponding to microtubule binding, ion channel binding, syntaxin binding and tubulin binding (Fig. 9B). Among genes more abundant in the hTN, suggesting strong axonal transport, we found transmembrane proteins like CRIM1, suggesting that local translation of some proteins may be performed efficiently near their site of action. Additionally we identified SCN5A and NTRK3, members of gene families known to be DRG-enriched, suggesting distinct molecular roles for members of these families in sensory neurons (NTRK1 abundances are enriched in hDRG over hTN) (Fig. 9C). The relative enrichment of all human genes between the hDRG and hTN nerve is detailed in Sheet 1 of Supplementary File 1.

## 4. Discussion

Our analysis profiled the hDRG transcriptome, and contrasted it with other human tissue transcriptomes and their corresponding mouse transcriptomes. By contrasting gene expression patterns across tissues in humans, we created comprehensive gene co-expression modules that take into account gene expression in DRGs. For each co-expression module, we identified putative transcription regulatory genes (TFs, and cofactors). For the co-expression module with genes specifically expressed in the hDRG (and silenced in other analyzed tissues), we identified 13 DRG-specific TFs. We confirmed known mammalian sensory development-related transcription factors in DRG like DRGX, TLX3 and PRDM12 in the adult hDRG, and additionally identified several transcription factors like POU4F1, POU4F3, HOXD1 and SKOR2. We comprehensively catalogued hDRG-expressed members of gene families known to be functionally important in the DRG: ion channels, receptors, kinases, and RNA binding proteins. We note that in the commonly used laboratory mouse strain, C57BL/6, DRG-specific genes are accurately predicted by hDRG-specificity suggesting strong translational capacity for this mouse model. Evolutionary conservation of DRG-specific gene expression suggests purifying selection, and supports the validity of the mouse as a preclinical model species for sensory biology and pain pharmacological and genetic research. Importantly, our work comprehensively characterized gene expression between hDRG and human tibial nerve, potentially identifying pervasive RNA transport to peripheral axons of hDRG neurons.

There are several factors that distinguish our work from previous studies that used high throughput transcriptome analyses. Our goal was to design an analysis framework to contrast the steady state mRNA profile of different human and mouse tissues, with a focus on identifying gene expression patterns across tissues that can be relevant for understanding DRG biology and guide pain therapeutic discovery. We focused on unbiased gene set enrichment analysis in DRG-specific gene expression patterns for functional analysis, in contrast to Flegel *et al* [25] who comprehensively characterized expression patterns of gene families known to be important in sensory tissues *a priori*. Flegel *et al* also pooled data from individual DRG donors *in vitro*, while we separately quantify transcriptome profiles from 3 individual DRG donors, pooling them *in silico* where required, such that both measures of dispersion and central tendency can be calculated on estimated transcript abundances for rigorous statistical analyses. Sapio *et al* [91] contrasted human and mouse DRG gene expression and performed functional analysis of DRG-enriched genes in the context of sensory neuropathies, but we additionally contrast mouse RNA-seq data from a homologous set of tissues to our human tissue panel, thereby building a framework to contrast human and mouse evolutionary divergence of expression patterns. Like the Sapio *et al* study, we identify constituent cell types where DRG-expressed genes are most likely to be transcribed. The most comprehensive study inclusive of the DRG for cross-tissue transcriptome comparisons between human and mouse was performed by the Genomics Institute of the Novartis Research Foundation in 2004 using standard Affymetrix microarrays [98]. To address these gaps and to generate our own resource for the field, we profiled the mRNA abundances in the hDRG by performing RNA-seq and contrasted it with relevant human and mouse RNA-seq datasets from public repositories, primarily the GTex Consortium [16]. Our quantification is based on a genome-wide readout of transcript abundance not limited to a predefined set of probes, has the added benefit of low technical variability with respect to microarray studies [55], and unlike the Novartis study is analyzed from the perspective of DRG biology, and drug target discovery in the PNS.

In addition to this, we performed several analyses which characterize the hDRG transcriptome in several biologically relevant ways for the first time. Over and above identifying hDRG-specific genes, gene co-expression analysis across human tissues allowed us to identify gene modules with similar co-expression patterns that are putatively co-regulated, thus taking the first step in identifying the transcriptional regulatory networks underlying fundamental cell types in the hDRG. Identification of hDRG-specific transcription factors allowed us to further identify transcription factors that may be implicated in controlling such regulatory networks into adulthood. Notably, our result divides the well-studied mammalian pan-CNS expressed genes [18] into CNS genes that are downregulated in the hDRG and those that are expressed in both the CNS and in the hDRG (such as DOCK3, which had been known to be expressed pan-CNS in mouse [71]).

Based on our comprehensively catalogued and annotated hDRG mRNA profile, we aim to perform several follow up studies. Since we specifically sourced L2 lumbar DRG from female donors, we aim to contrast mRNA profiles between both male and female DRG donors, to identify sex-specific differences in DRG gene expression. We aim to identify if there are region specific differences in the DRG transcriptome by analyzing DRGs from additional spinal levels and that innervate different target tissues. Finally, we aim to perform RNA-seq on cohorts of pain patients to identify both patient-specific and cohort-specific transcriptomic signatures of chronic pain, if they exist, in the hDRG. The absence of these datasets in the present analysis can be viewed as a weakness of this study, but, to our knowledge, these datasets do not exist or are not publicly available. All of our sequencing data is publicly available and can be searched without the need for additional analysis on our resource webpage. This resource can be built upon by interested researchers.

A primary novel finding of our work is evidence for pervasive trafficking of RNA into the peripheral axons of DRG neurons. Comparing L2 transcriptomes from our own RNA-seq experiments to similar experiments conducted by the GTex Consortium on human tibial nerve samples suggests a dramatic overlap in transcriptomes that cannot be accounted for simply by Schwann cell or fibroblast transcriptomes. We interpret this to mean that RNAs generated in DRG neuron nuclei are trafficked into peripherally projecting axons, likely through binding to RNA binding proteins that have previously been shown to be abundant in DRG axons [83; 84]. This finding opens the possibility of gaining insight into changes in DRG transcriptomes in humans by sampling from peripheral nerves (e.g. tibial nerve biopsies) or even from skin biopsies that are routinely taken for evaluation of epidermal nerve fiber densities in neuropathic pain patients. By comparing transcriptomes of peripheral tissues like skin biopsies with nerve endings, or nerve biopsies, we aim to eventually be able to build up a “personal genomics” model of the pain transcriptome [108] that can give molecular neurobiological insight into phenotype changes in DRG neurons that drive chronic pain. Insofar as local translation at nociceptor terminals has already been identified as a key regulation step in nociceptor plasticity [58–60], this insight could develop into a powerful tool for mechanism-based discovery in patients.

## Acknowledgements

This work was supported by NIH grants R01NS065926 (TJP), R01GM102575 (TJP and GD), R01MH102616 and R01MH109665 (MQZ) and The University of Texas STARS program (TJP, GD and MZ).

## Author Contributions

TJP conceived the project. TJP and GD guided research and supervised all personnel. TJP and PR designed RNA sequencing study. PR conceived and designed all bioinformatics studies. PR implemented all bioinformatics analyses with the following inputs: AT implemented the RNA-seq pipeline, LQ implemented the microarray pipeline, and AW implemented hierarchical clustering of assays. MN, CR, TL and PR coded the companion web service. J-YK, CR, MN, AW and PR performed pilot experiments. THK and MQZ provided feedback about RNA-seq computational analysis. PR drew all figures. PR and TJP wrote the paper. The authors wish to thank Anagha Krishnan and Yashas Manjunatha for data collation.

## Supplementary figures

**Supplementary Figure 1.**
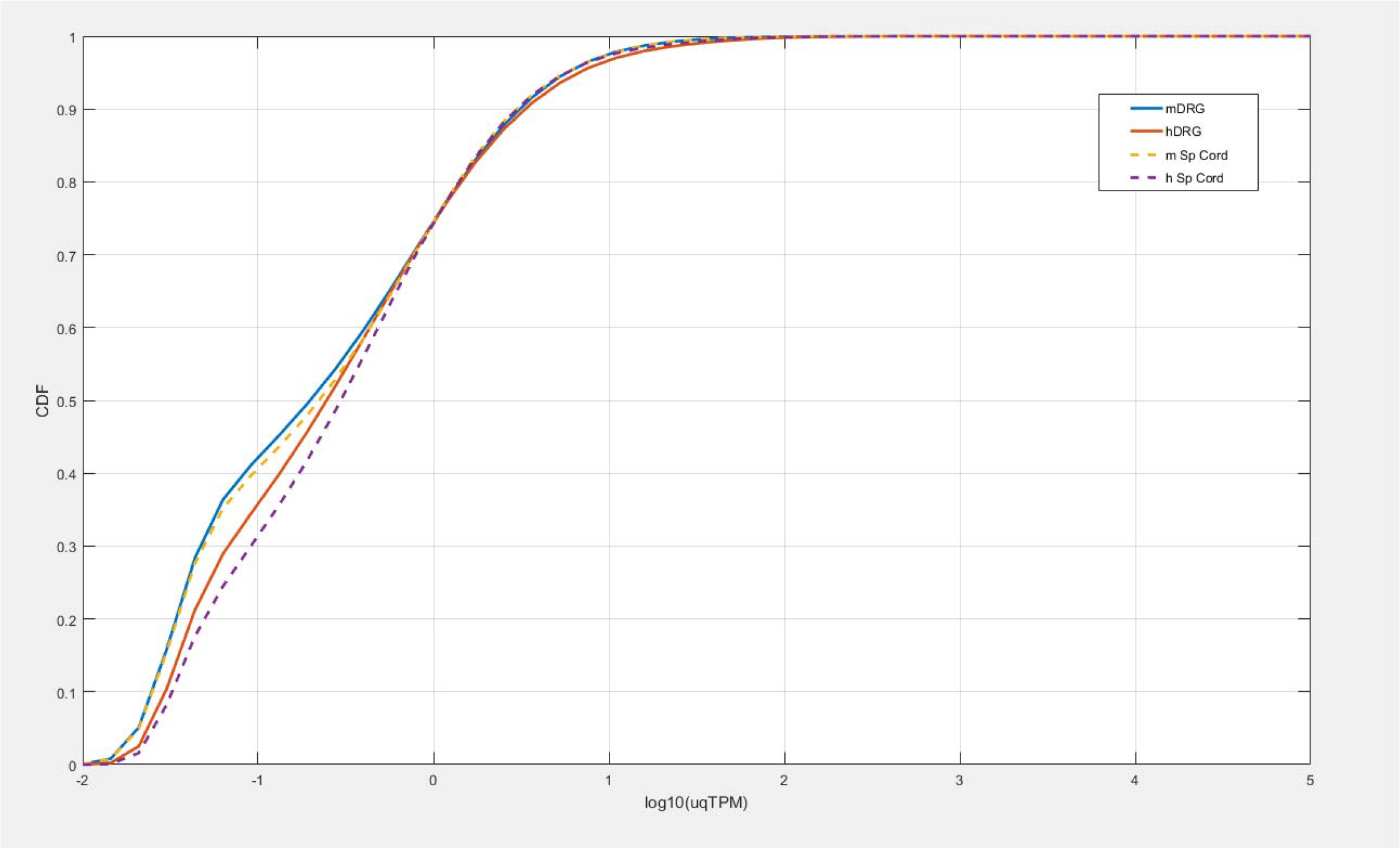
Cumulative distribution function for human and mouse DRG and spinal cord relative abundances in uqTPM. The CDFs track in approximately similar ways above 0 uqTPM. Differences at lower uqTPMs can potentially be attributed to differing sequencing depths.

**Supplementary Figure 2.**
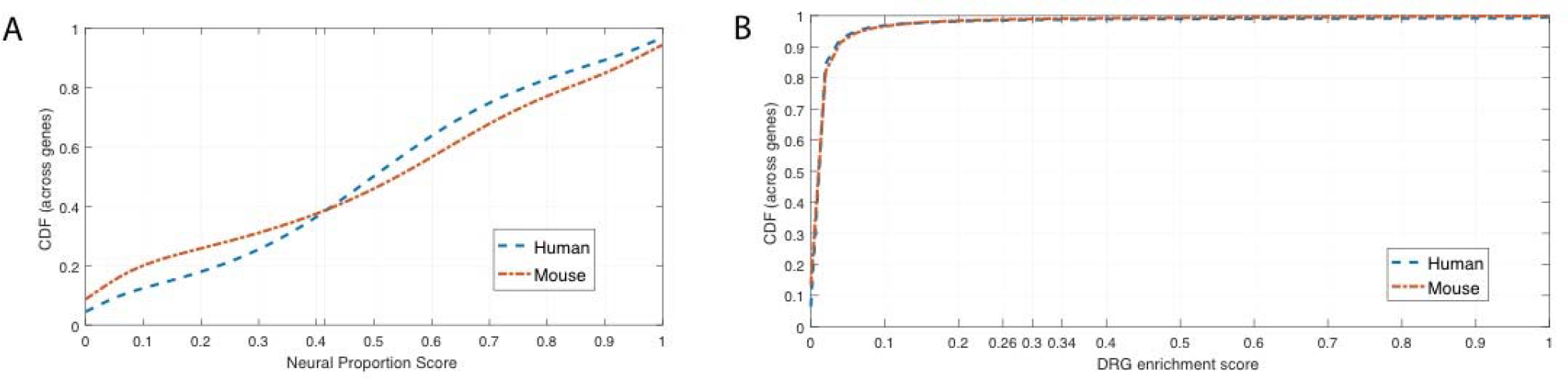
Cumulative distribution function for **(A)** human and mouse neural proportion scores for coding genes **(B)** human and mouse DRG enrichment scores for coding genes. CDFs for DRG enrichment scores are similar, suggesting a similar number of genes are DRG-enriched in human and mouse. The CDFs for neural proportion scores differ between human and mouse, possibly due to the evolutionary divergence of the CNS transcriptome between human and mouse.

**Supplementary Figure 3.**
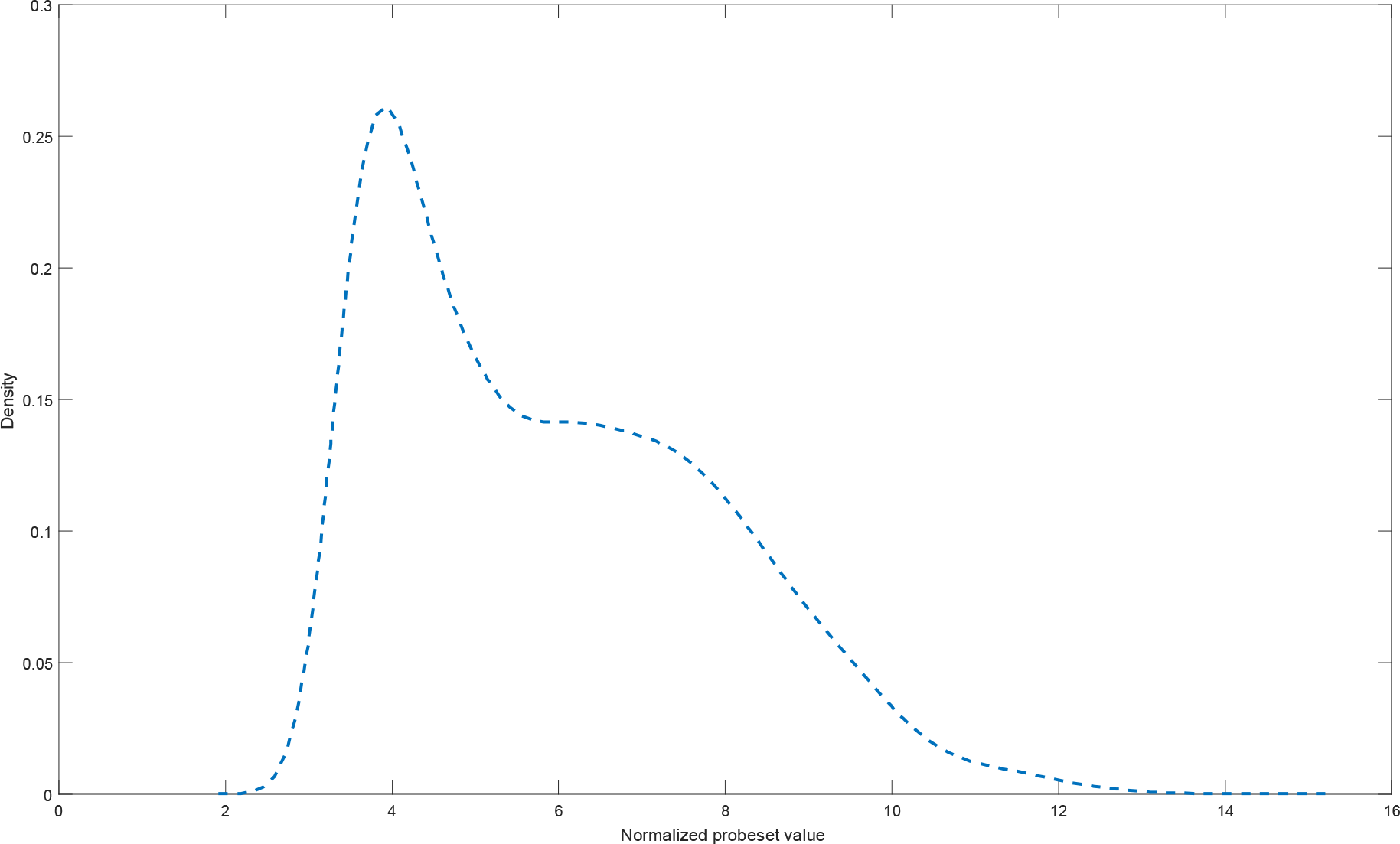
Estimated density function for normalized probeset values for analyzed microarray datasets for human DRG, TG, cultured NHSCs, and cultured fibroblasts.

**Supplementary Figure 4.**
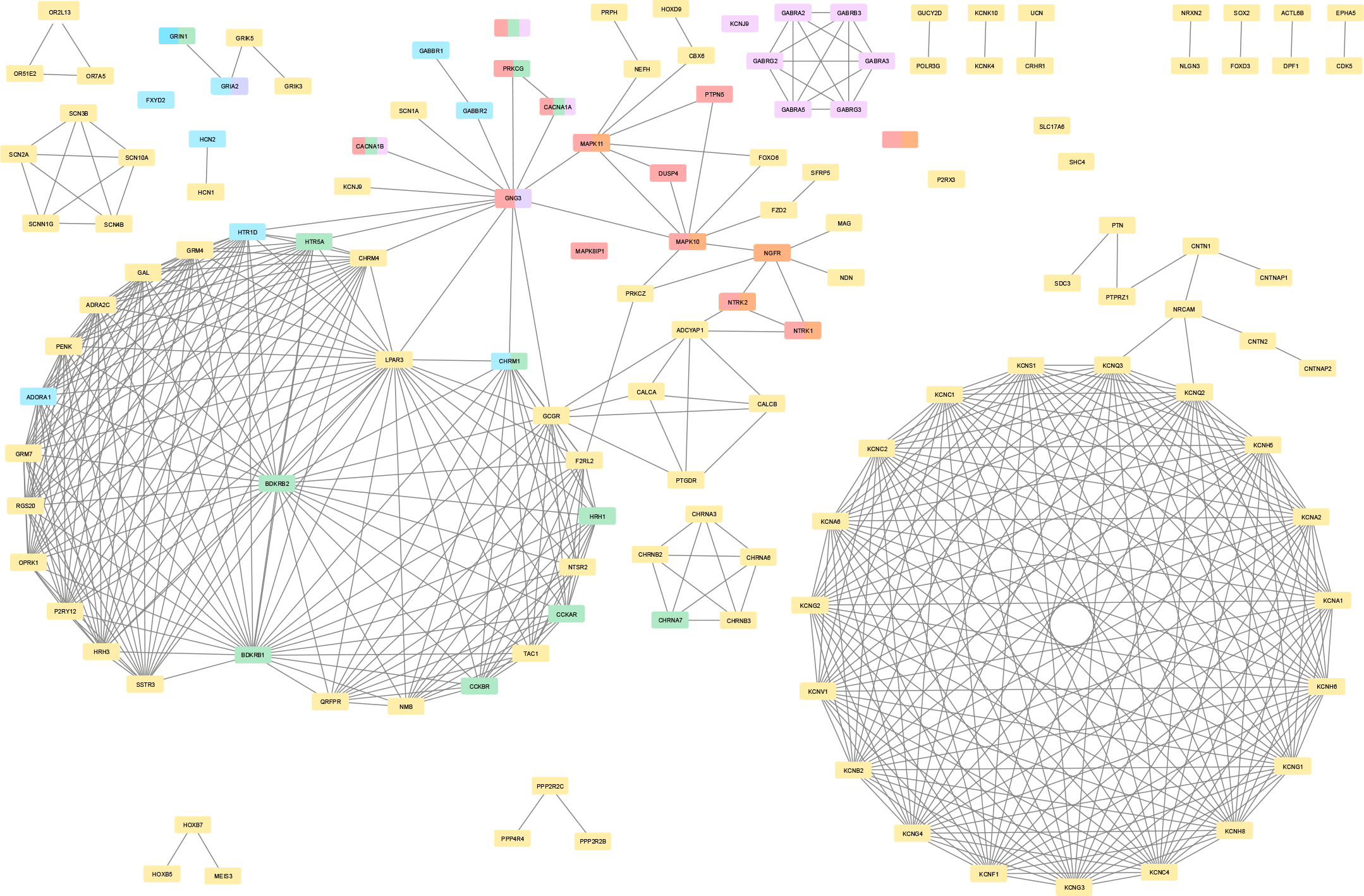
PPI subnetwork constructed from StringDB database for genes that are strongly hDRG-enriched or weakly hDRG-enriched and a member of one of the profiled gene families we analyzed. We identified several pathways that overlap the genes involved in our reconstructed PPI subnetwork.

## Notes

**Conflict of interest:** The authors declare no conflicts of interest

## References

[1] Abaffy T. Human Olfactory Receptors Expression and Their Role in Non-Olfactory Tissues-A Mini-Review. Journal of Pharmacogenomics & Pharmacoproteomics 2015;6(4):1.

[2] Ambjørn M, Dubreuil V, Miozzo F, Nigon F, Møller B, Issazadeh-Navikas S, Berg J, Lees M, Sap J. A loss-of-function screen for phosphatases that regulate neurite outgrowth identifies PTPN12 as a negative regulator of TrkB tyrosine phosphorylation. PloS one 2013;8(6):e65371.

[3] Ashburner M, Ball CA, Blake JA, Botstein D, Butler H, Cherry JM, Davis AP, Dolinski K, Dwight SS, Eppig JT, Harris MA, Hill DP, Issel-Tarver L, Kasarskis A, Lewis S, Matese JC, Richardson JE, Ringwald M, Rubin GM, Sherlock G. Gene ontology: tool for the unification of biology. The Gene Ontology Consortium. Nat Genet 2000;25(1):25–29.

[4] Atluri S, Sudarshan G, Manchikanti L. Assessment of the trends in medical use and misuse of opioid analgesics from 2004 to 2011. Pain Physician 2014;17(2):E119–128.

[5] Bannwarth B, Kostine M. Targeting nerve growth factor (NGF) for pain management: what does the future hold for NGF antagonists? Drugs 2014;74(6):619–626.

[6] Bar-Joseph Z, Gifford DK, Jaakkola TS. Fast optimal leaf ordering for hierarchical clustering. Bioinformatics 2001;17(suppl 1):S22–S29.

[7] Blanchard JW, Eade KT, Szűcs A, Sardo VL, Tsunemoto RK, Williams D, Sanna PP, Baldwin KK. Selective conversion of fibroblasts into peripheral sensory neurons. Nature neuroscience 2015;18(1):25–35.

[8] Bolstad BM, Irizarry RA, Åstrand M, Speed TP. A comparison of normalization methods for high density oligonucleotide array data based on variance and bias. Bioinformatics 2003;19(2):185–193.

[9] Carvalho BS, Irizarry RA. A framework for oligonucleotide microarray preprocessing. Bioinformatics 2010;26(19):2363–2367.

[10] Chambers SM, Qi Y, Mica Y, Lee G, Zhang X-J, Niu L, Bilsland J, Cao L, Stevens E, Whiting P. Combined small-molecule inhibition accelerates developmental timing and converts human pluripotent stem cells into nociceptors. Nature biotechnology 2012;30(7):715–720.

[11] Chen EY, Tan CM, Kou Y, Duan Q, Wang Z, Meirelles GV, Clark NR, Ma’ayan A. Enrichr: interactive and collaborative HTML5 gene list enrichment analysis tool. BMC Bioinformatics 2013;14:128.

[12] Chen YC, Auer-Grumbach M, Matsukawa S, Zitzelsberger M, Themistocleous AC, Strom TM, Samara C, Moore AW, Cho LT, Young GT, Weiss C, Schabhuttl M, Stucka R, Schmid AB, Parman Y, Graul-Neumann L, Heinritz W, Passarge E, Watson RM, Hertz JM, Moog U, Baumgartner M, Valente EM, Pereira D, Restrepo CM, Katona I, Dusl M, Stendel C, Wieland T, Stafford F, Reimann F, von Au K, Finke C, Willems PJ, Nahorski MS, Shaikh SS, Carvalho OP, Nicholas AK, Karbani G, McAleer MA, Cilio MR, McHugh JC, Murphy SM, Irvine AD, Jensen UB, Windhager R, Weis J, Bergmann C, Rautenstrauss B, Baets J, De Jonghe P, Reilly MM, Kropatsch R, Kurth I, Chrast R, Michiue T, Bennett DL, Woods CG, Senderek J. Transcriptional regulator PRDM12 is essential for human pain perception. Nat Genet 2015;47(7):803–808.

[13] Chinwalla AT, Cook LL, Delehaunty KD, Fewell GA, Fulton LA, Fulton RS, Graves TA, Hillier LW, Mardis ER, McPherson JD. Initial sequencing and comparative analysis of the mouse genome. Nature 2002;420(6915):520–562.

[14] Chiu IM, Barrett LB, Williams EK, Strochlic DE, Lee S, Weyer AD, Lou S, Bryman GS, Roberson DP, Ghasemlou N, Piccoli C, Ahat E, Wang V, Cobos EJ, Stucky CL, Ma Q, Liberles SD, Woolf CJ. Transcriptional profiling at whole population and single cell levels reveals somatosensory neuron molecular diversity. Elife 2014;3.

[15] Consortium GO. Gene Ontology annotations and resources. Nucleic acids research 2013;41(D1):D530–D535.

[16] Consortium GT. The Genotype-Tissue Expression (GTEx) project. Nat Genet 2013;45(6):580–585.

[17] Croft D, Mundo AF, Haw R, Milacic M, Weiser J, Wu G, Caudy M, Garapati P, Gillespie M, Kamdar MR, Jassal B, Jupe S, Matthews L, May B, Palatnik S, Rothfels K, Shamovsky V, Song H, Williams M, Birney E, Hermjakob H, Stein L, D’Eustachio P. The Reactome pathway knowledgebase. Nucleic Acids Res 2014;42(Database issue):D472–477.

[18] Darmanis S, Sloan SA, Zhang Y, Enge M, Caneda C, Shuer LM, Gephart MGH, Barres BA, Quake SR. A survey of human brain transcriptome diversity at the single cell level. Proceedings of the National Academy of Sciences 2015;112(23):7285–7290.

[19] Dart RC, Surratt HL, Cicero TJ, Parrino MW, Severtson SG, Bucher-Bartelson B, Green JL. Trends in opioid analgesic abuse and mortality in the United States. N Engl J Med 2015;372(3):241–248.

[20] Dawes JM, Antunes-Martins A, Perkins JR, Paterson KJ, Sisignano M, Schmid R, Rust W, Hildebrandt T, Geisslinger G, Orengo C, Bennett DL, McMahon SB. Genome-wide transcriptional profiling of skin and dorsal root ganglia after ultraviolet-B-induced inflammation. PLoS One 2014;9(4):e93338.

[21] Dreszer TR, Karolchik D, Zweig AS, Hinrichs AS, Raney BJ, Kuhn RM, Meyer LR, Wong M, Sloan CA, Rosenbloom KR, Roe G, Rhead B, Pohl A, Malladi VS, Li CH, Learned K, Kirkup V, Hsu F, Harte RA, Guruvadoo L, Goldman M, Giardine BM, Fujita PA, Diekhans M, Cline MS, Clawson H, Barber GP, Haussler D, James Kent W. The UCSC Genome Browser database: extensions and updates 2011. Nucleic Acids Res 2012;40(Database issue):D918–923.

[22] Egleton RD, Davis TP. Development of neuropeptide drugs that cross the blood-brain barrier. NeuroRx 2005;2(1):44–53.

[23] Eisen MB, Spellman PT, Brown PO, Botstein D. Cluster analysis and display of genome-wide expression patterns. Proc Natl Acad Sci U S A 1998;95(25):14863–14868.

[24] Felfly H, Muotri A, Yao H, Haddad GG. Hematopoietic stem cell transplantation protects mice from lethal stroke. Exp Neurol 2010;225(2):284–293.

[25] Flegel C, Schobel N, Altmuller J, Becker C, Tannapfel A, Hatt H, Gisselmann G. RNA-Seq Analysis of Human Trigeminal and Dorsal Root Ganglia with a Focus on Chemoreceptors. PLoS One 2015;10(6):e0128951.

[26] Franceschini A. STRINGdb Package Vignette. Nucleic Acids Res 2013.

[27] Geer LY, Marchler-Bauer A, Geer RC, Han L, He J, He S, Liu C, Shi W, Bryant SH. The NCBI BioSystems database. Nucleic Acids Res 2010;38(Database issue):D492–496.

[28] Glusman G, Caballero J, Robinson M, Kutlu B, Hood L. Optimal scaling of digital transcriptomes. PloS one 2013;8(11):e77885.

[29] Goswami SC, Mishra SK, Maric D, Kaszas K, Gonnella GL, Clokie SJ, Kominsky HD, Gross JR, Keller JM, Mannes AJ, Hoon MA, Iadarola MJ. Molecular signatures of mouse TRPV1-lineage neurons revealed by RNA-Seq transcriptome analysis. J Pain 2014;15(12):1338–1359.

[30] Gray KA, Seal RL, Tweedie S, Wright MW, Bruford EA. A review of the new HGNC gene family resource. Human genomics 2016;10(1):6.

[31] Griebel G, Holsboer F. Neuropeptide receptor ligands as drugs for psychiatric diseases: the end of the beginning? Nature Reviews Drug Discovery 2012;11(6):462–478.

[32] Guan Z, Kuhn JA, Wang X, Colquitt B, Solorzano C, Vaman S, Guan AK, Evans-Reinsch Z, Braz J, Devor M, Abboud-Werner SL, Lanier LL, Lomvardas S, Basbaum AI. Injured sensory neuron-derived CSF1 induces microglial proliferation and DAP12-dependent pain. Nat Neurosci 2016;19(1):94–101.

[33] Harrow J, Frankish A, Gonzalez JM, Tapanari E, Diekhans M, Kokocinski F, Aken BL, Barrell D, Zadissa A, Searle S, Barnes I, Bignell A, Boychenko V, Hunt T, Kay M, Mukherjee G, Rajan J, Despacio-Reyes G, Saunders G, Steward C, Harte R, Lin M, Howald C, Tanzer A, Derrien T, Chrast J, Walters N, Balasubramanian S, Pei B, Tress M, Rodriguez JM, Ezkurdia I, van Baren J, Brent M, Haussler D, Kellis M, Valencia A, Reymond A, Gerstein M, Guigo R, Hubbard TJ. GENCODE: the reference human genome annotation for The ENCODE Project. Genome Res 2012;22(9):1760–1774.

[34] Hill MO. Diversity and evenness: a unifying notation and its consequences. Ecology 1973;54(2):427–432.

[35] Hu G, Huang K, Hu Y, Du G, Xue Z, Zhu X, Fan G. Single-cell RNA-seq reveals distinct injury responses in different types of DRG sensory neurons. Sci Rep 2016;6:31851.

[36] Inquimbert P, Bartels K, Babaniyi OB, Barrett LB, Tegeder I, Scholz J. Peripheral nerve injury produces a sustained shift in the balance between glutamate release and uptake in the dorsal horn of the spinal cord. Pain 2012;153(12):2422–2431.

[37] Isensee J, Wenzel C, Buschow R, Weissmann R, Kuss AW, Hucho T. Subgroup-elimination transcriptomics identifies signaling proteins that define subclasses of TRPV1-positive neurons and a novel paracrine circuit. PLoS One 2014;9(12):e115731.

[38] Ji SJ, Jaffrey SR. Axonal transcription factors: Novel regulators of growth cone-to-nucleus signaling. Developmental neurobiology 2014;74(3):245–258.

[39] Jiang N, Li H, Sun Y, Yin D, Zhao Q, Cui S, Yao D. Differential gene expression in proximal and distal nerve segments of rats with sciatic nerve injury during Wallerian degeneration. Neural Regen Res 2014;9(12):1186–1194.

[40] Jiang Y, Clark WT, Friedberg I, Radivojac P. The impact of incomplete knowledge on the evaluation of protein function prediction: a structured-output learning perspective. Bioinformatics 2014;30(17):i609–i616.

[41] Jung H, Yoon BC, Holt CE. Axonal mRNA localization and local protein synthesis in nervous system assembly, maintenance and repair. Nat Rev Neurosci 2012;13(5):308–324.

[42] Kanehisa M, Sato Y, Kawashima M, Furumichi M, Tanabe M. KEGG as a reference resource for gene and protein annotation. Nucleic acids research 2015:gkv1070.

[43] Kim AY, Tang Z, Liu Q, Patel KN, Maag D, Geng Y, Dong X. Pirt, a phosphoinositide-binding protein, functions as a regulatory subunit of TRPV1. Cell 2008;133(3):475–485.

[44] Kim YS, Chu Y, Han L, Li M, Li Z, LaVinka PC, Sun S, Tang Z, Park K, Caterina MJ. Central terminal sensitization of TRPV1 by descending serotonergic facilitation modulates chronic pain. Neuron 2014;81(4):873–887.

[45] Kiryushko D, Bock E, Berezin V. Pharmacology of cell adhesion molecules of the nervous system. Current neuropharmacology 2007;5(4):253–267.

[46] Kogelman LJA, Christensen RE, Pedersen SH, Bertalan M, Hansen TF, Jansen-Olesen I, Olesen J. Whole transcriptome expression of trigeminal ganglia compared to dorsal root ganglia in Rattus Norvegicus. Neuroscience 2017;350:169–179.

[47] Kuja-Panula J, Kiiltomäki M, Yamashiro T, Rouhiainen A, Rauvala H. AMIGO, a transmembrane protein implicated in axon tract development, defines a novel protein family with leucine-rich repeats. The Journal of cell biology 2003;160(6):963–973.

[48] Kuleshov MV, Jones MR, Rouillard AD, Fernandez NF, Duan Q, Wang Z, Koplev S, Jenkins SL, Jagodnik KM, Lachmann A, McDermott MG, Monteiro CD, Gundersen GW, Ma’ayan A. Enrichr: a comprehensive gene set enrichment analysis web server 2016 update. Nucleic Acids Res 2016;44(W1):W90–97.

[49] Kupczyk P. Expression Of UCHL1 Gene In The Skin Of Psoriasis: Neuroepidermal Marker Of Itch.

[50] Kutuzov MA, Solov’eva OV, Andreeva AV, Bennett N. Protein Ser/Thr phosphatases PPEF interact with calmodulin. Biochemical and biophysical research communications 2002;293(3):1047–1052.

[51] Li CL, Li KC, Wu D, Chen Y, Luo H, Zhao JR, Wang SS, Sun MM, Lu YJ, Zhong YQ, Hu XY, Hou R, Zhou BB, Bao L, Xiao HS, Zhang X. Somatosensory neuron types identified by high-coverage single-cell RNA-sequencing and functional heterogeneity. Cell Res 2016;26(1):83–102.

[52] Liu Q, Dong X. The role of the Mrgpr receptor family in itch. Pharmacology of Itch: Springer, 2015. pp. 71–88.

[53] Lopes C, Liu Z, Xu Y, Ma Q. Tlx3 and Runx1 act in combination to coordinate the development of a cohort of nociceptors, thermoceptors, and pruriceptors. J Neurosci 2012;32(28):9706–9715.

[54] Manteniotis S, Lehmann R, Flegel C, Vogel F, Hofreuter A, Schreiner BS, Altmuller J, Becker C, Schobel N, Hatt H, Gisselmann G. Comprehensive RNA-Seq expression analysis of sensory ganglia with a focus on ion channels and GPCRs in Trigeminal ganglia. PLoS One 2013;8(11):e79523.

[55] Marioni JC, Mason CE, Mane SM, Stephens M, Gilad Y. RNA-seq: an assessment of technical reproducibility and comparison with gene expression arrays. Genome Res 2008;18(9):1509–1517.

[56] Masahira N, Takebayashi H, Ono K, Watanabe K, Ding L, Furusho M, Ogawa Y, Nabeshima Y, Alvarez-Buylla A, Shimizu K, Ikenaka K. Olig2-positive progenitors in the embryonic spinal cord give rise not only to motoneurons and oligodendrocytes, but also to a subset of astrocytes and ependymal cells. Dev Biol 2006;293(2):358–369.

[57] McCleane GJ, Suzuki R, Dickenson AH. Does a single intravenous injection of the 5HT3 receptor antagonist ondansetron have an analgesic effect in neuropathic pain? A double-blinded, placebo-controlled cross-over study. Anesthesia & Analgesia 2003;97(5):1474–1478.

[58] Melemedjian OK, Asiedu MN, Tillu DV, Peebles KA, Yan J, Ertz N, Dussor GO, Price TJ. IL-6- and NGF-induced rapid control of protein synthesis and nociceptive plasticity via convergent signaling to the eIF4F complex. J Neurosci 2010;30(45):15113–15123.

[59] Melemedjian OK, Asiedu MN, Tillu DV, Sanoja R, Yan J, Lark A, Khoutorsky A, Johnson J, Peebles KA, Lepow T, Sonenberg N, Dussor G, Price TJ. Targeting adenosine monophosphate-activated protein kinase (AMPK) in preclinical models reveals a potential mechanism for the treatment of neuropathic pain. Mol Pain 2011;7:70.

[60] Melemedjian OK, Tillu DV, Moy JK, Asiedu MN, Mandell EK, Ghosh S, Dussor G, Price TJ. Local translation and retrograde axonal transport of CREB regulates IL-6-induced nociceptive plasticity. Mol Pain 2014;10:45.

[61] Miller JA, Horvath S, Geschwind DH. Divergence of human and mouse brain transcriptome highlights Alzheimer disease pathways. Proc Natl Acad Sci U S A 2010;107(28):12698–12703.

[62] Miller SJ, Jessen WJ, Mehta T, Hardiman A, Sites E, Kaiser S, Jegga AG, Li H, Upadhyaya M, Giovannini M, Muir D, Wallace MR, Lopez E, Serra E, Nielsen GP, Lazaro C, Stemmer-Rachamimov A, Page G, Aronow BJ, Ratner N. Integrative genomic analyses of neurofibromatosis tumours identify SOX9 as a biomarker and survival gene. EMBO Mol Med 2009;1(4):236–248.

[63] Minis A, Dahary D, Manor O, Leshkowitz D, Pilpel Y, Yaron A. Subcellular transcriptomics-dissection of the mRNA composition in the axonal compartment of sensory neurons. Dev Neurobiol 2014;74(3):365–381.

[64] Mishra SK, Holzman S, Hoon MA. A nociceptive signaling role for neuromedin B. Journal of Neuroscience 2012;32(25):8686–8695.

[65] Molyneaux BJ, Arlotta P, Menezes JR, Macklis JD. Neuronal subtype specification in the cerebral cortex. Nat Rev Neurosci 2007;8(6):427–437.

[66] Momin A, Wood JN. Sensory neuron voltage-gated sodium channels as analgesic drug targets. Curr Opin Neurobiol 2008;18(4):383–388.

[67] Moore DL, Goldberg JL. Multiple transcription factor families regulate axon growth and regeneration. Developmental neurobiology 2011;71(12):1186–1211.

[68] Nadjar Y, Triller A, Bessereau J-L, Dumoulin A. The Susd2 protein regulates neurite growth and excitatory synaptic density in hippocampal cultures. Molecular and Cellular Neuroscience 2015;65:82–91.

[69] Nagy V, Cole T, Van Campenhout C, Khoung TM, Leung C, Vermeiren S, Novatchkova M, Wenzel D, Cikes D, Polyansky AA, Kozieradzki I, Meixner A, Bellefroid EJ, Neely GG, Penninger JM. The evolutionarily conserved transcription factor PRDM12 controls sensory neuron development and pain perception. Cell Cycle 2015;14(12):1799–1808.

[70] Nakatani T, Minaki Y, Kumai M, Nitta C, Ono Y. The c-Ski family member and transcriptional regulator Corl2/Skor2 promotes early differentiation of cerebellar Purkinje cells. Developmental biology 2014;388(1):68–80.

[71] Namekata K, Enokido Y, Iwasawa K, Kimura H. MOCA induces membrane spreading by activating Rac1. Journal of Biological Chemistry 2004;279(14):14331–14337.

[72] Nascimento R, Santiago M, Marques S, Allodi S, Martinez A. Diversity among satellite glial cells in dorsal root ganglia of the rat. Brazilian Journal of Medical and Biological Research 2008;41(11):1011–1017.

[73] Niederreither K, Vermot J, Schuhbaur B, Chambon P, Dolle P. Retinoic acid synthesis and hindbrain patterning in the mouse embryo. Development 2000;127(1):75–85.

[74] Nomaksteinsky M, Kassabov S, Chettouh Z, Stoekle HC, Bonnaud L, Fortin G, Kandel ER, Brunet JF. Ancient origin of somatic and visceral neurons. BMC Biol 2013;11:53.

[75] Okaty BW, Sugino K, Nelson SB. Cell type-specific transcriptomics in the brain. Journal of Neuroscience 2011;31(19):6939–6943.

[76] Oldham MC, Horvath S, Geschwind DH. Conservation and evolution of gene coexpression networks in human and chimpanzee brains. Proc Natl Acad Sci U S A 2006;103(47):17973–17978.

[77] Oort PJ, Warden CH, Baumann TK, Knotts TA, Adams SH. Characterization of Tusc5, an adipocyte gene co-expressed in peripheral neurons. Molecular and cellular endocrinology 2007;276(1):24–35.

[78] Owens ND, Blitz IL, Lane MA, Patrushev I, Overton JD, Gilchrist MJ, Cho KW, Khokha MK. Measuring absolute RNA copy numbers at high temporal resolution reveals transcriptome kinetics in development. Cell reports 2016;14(3):632–647.

[79] Pachter L. Models for transcript quantification from RNA-Seq. arXiv preprint arXiv:11043889 2011.

[80] Pacini A, Micheli L, Maresca M, Branca JJV, McIntosh JM, Ghelardini C, Mannelli LDC. The α9α10 nicotinic receptor antagonist a-conotoxin RgIA prevents neuropathic pain induced by oxaliplatin treatment. Experimental neurology 2016;282:37–48.

[81] Pearson K. Note on regression and inheritance in the case of two parents. Proceedings of the Royal Society of London 1895;58:240–242.

[82] Pevny LH, Sockanathan S, Placzek M, Lovell-Badge R. A role for SOX1 in neural determination. Development 1998;125(10):1967–1978.

[83] Price TJ, Hargreaves KM, Cervero F. Protein expression and mRNA cellular distribution of the NKCC1 cotransporter in the dorsal root and trigeminal ganglia of the rat. Brain research 2006;1112(1):146–158.

[84] Price TJ, Rashid MH, Millecamps M, Sanoja R, Entrena JM, Cervero F. Decreased nociceptive sensitization in mice lacking the fragile X mental retardation protein: role of mGluR1/5 and mTOR. Journal of Neuroscience 2007;27(51):13958–13967.

[85] Ramskold D, Wang ET, Burge CB, Sandberg R. An abundance of ubiquitously expressed genes revealed by tissue transcriptome sequence data. PLoS Comput Biol 2009;5(12):e1000598.

[86] Rebelo S, Chen ZF, Anderson DJ, Lima D. Involvement of DRG11 in the development of the primary afferent nociceptive system. Mol Cell Neurosci 2006;33(3):236–246.

[87] Romero IG, Pai AA, Tung J, Gilad Y. RNA-seq: impact of RNA degradation on transcript quantification. BMC biology 2014;12(1):42.

[88] Roth RB, Hevezi P, Lee J, Willhite D, Lechner SM, Foster AC, Zlotnik A. Gene expression analyses reveal molecular relationships among 20 regions of the human CNS. Neurogenetics 2006;7(2):67–80.

[89] Rouwette T, Sondermann J, Avenali L, Gomez-Varela D, Schmidt M. Standardized Profiling of The Membrane-Enriched Proteome of Mouse Dorsal Root Ganglia (DRG) Provides Novel Insights Into Chronic Pain. Mol Cell Proteomics 2016;15(6):2152–2168.

[90] Roux J, Rosikiewicz M, Robinson-Rechavi M. What to compare and how: Comparative transcriptomics for Evo-Devo. Journal of Experimental Zoology Part B: Molecular and Developmental Evolution 2015;324(4):372–382.

[91] Sapio MR, Goswami SC, Gross JR, Mannes AJ, Iadarola MJ. Transcriptomic analyses of genes and tissues in inherited sensory neuropathies. Exp Neurol 2016;283(Pt A):375–395.

[92] Scholz J, Woolf CJ. Can we conquer pain? Nat Neurosci 2002;5 Suppl:1062–1067.

[93] Schwertassek U, Buckley DA, Xu CF, Lindsay AJ, McCaffrey MW, Neubert TA, Tonks NK. Myristoylation of the dual-specificity phosphatase c-JUN N-terminal kinase (JNK) stimulatory phosphatase 1 is necessary for its activation of JNK signaling and apoptosis. Febs Journal 2010;277(11):2463–2473.

[94] Seok J, Warren HS, Cuenca AG, Mindrinos MN, Baker HV, Xu W, Richards DR, McDonald-Smith GP, Gao H, Hennessy L, Finnerty CC, Lopez CM, Honari S, Moore EE, Minei JP, Cuschieri J, Bankey PE, Johnson JL, Sperry J, Nathens AB, Billiar TR, West MA, Jeschke MG, Klein MB, Gamelli RL, Gibran NS, Brownstein BH, Miller-Graziano C, Calvano SE, Mason PH, Cobb JP, Rahme LG, Lowry SF, Maier RV, Moldawer LL, Herndon DN, Davis RW, Xiao W, Tompkins RG, Inflammation, Host Response to Injury LSCRP. Genomic responses in mouse models poorly mimic human inflammatory diseases. Proc Natl Acad Sci U S A 2013;110(9):3507–3512.

[95] Shannon CE. Communication in the presence of noise. Proceedings of the IRE 1949;37(1):10–21.

[96] Shiroguchi K, Jia TZ, Sims PA, Xie XS. Digital RNA sequencing minimizes sequence-dependent bias and amplification noise with optimized single-molecule barcodes. Proceedings of the National Academy of Sciences 2012;109(4):1347–1352.

[97] Short KM, Cox TC. Subclassification of the RBCC/TRIM superfamily reveals a novel motif necessary for microtubule binding. Journal of Biological Chemistry 2006;281(13):8970–8980.

[98] Su AI, Wiltshire T, Batalov S, Lapp H, Ching KA, Block D, Zhang J, Soden R, Hayakawa M, Kreiman G, Cooke MP, Walker JR, Hogenesch JB. A gene atlas of the mouse and human protein-encoding transcriptomes. Proc Natl Acad Sci U S A 2004;101(16):6062–6067.

[99] Sudmant PH, Alexis MS, Burge CB. Meta-analysis of RNA-seq expression data across species, tissues and studies. Genome Biol 2015;16:287.

[100] Takao K, Miyakawa T. Genomic responses in mouse models greatly mimic human inflammatory diseases. Proceedings of the National Academy of Sciences 2015;112(4):1167–1172.

[101] Tang Z, Kim A, Masuch T, Park K, Weng H, Wetzel C, Dong X. Pirt functions as an endogenous regulator of TRPM8. Nat Commun 2013;4:2179.

[102] Thakur M, Crow M, Richards N, Davey GI, Levine E, Kelleher JH, Agley CC, Denk F, Harridge SD, McMahon SB. Defining the nociceptor transcriptome. Front Mol Neurosci 2014;7:87.

[103] Trapnell C, Hendrickson DG, Sauvageau M, Goff L, Rinn JL, Pachter L. Differential analysis of gene regulation at transcript resolution with RNA-seq. Nat Biotechnol 2013;31(1):46–53.

[104] Trapnell C, Pachter L, Salzberg SL. TopHat: discovering splice junctions with RNA-Seq. Bioinformatics 2009;25(9):1105–1111.

[105] Trapnell C, Roberts A, Goff L, Pertea G, Kim D, Kelley DR, Pimentel H, Salzberg SL, Rinn JL, Pachter L. Differential gene and transcript expression analysis of RNA-seq experiments with TopHat and Cufflinks. Nat Protoc 2012;7(3):562–578.

[106] Usoskin D, Furlan A, Islam S, Abdo H, Lönnerberg P, Lou D, Hjerling-Leffler J, Haeggström J, Kharchenko O, Kharchenko PV. Unbiased classification of sensory neuron types by large-scale single-cell RNA sequencing. Nature neuroscience 2015;18(1):145–153.

[107] Usoskin D, Furlan A, Islam S, Abdo H, Lonnerberg P, Lou D, Hjerling-Leffler J, Haeggstrom J, Kharchenko O, Kharchenko PV, Linnarsson S, Ernfors P. Unbiased classification of sensory neuron types by large-scale single-cell RNA sequencing. Nat Neurosci 2015;18(1):145–153.

[108] Vellucci R. Heterogeneity of chronic pain. Clin Drug Investig 2012;32 Suppl 1:3–10.

[109] Vilella AJ, Severin J, Ureta-Vidal A, Heng L, Durbin R, Birney E. EnsemblCompara GeneTrees: Complete, duplication-aware phylogenetic trees in vertebrates. Genome research 2009;19(2):327–335.

[110] Wagner AH, Coffman AC, Ainscough BJ, Spies NC, Skidmore ZL, Campbell KM, Krysiak K, Pan D, McMichael JF, Eldred JM, Walker JR, Wilson RK, Mardis ER, Griffith M, Griffith OL. DGIdb 2.0: mining clinically relevant drug-gene interactions. Nucleic Acids Res 2016;44(D1):D1036–1044.

[111] Wainger BJ, Buttermore ED, Oliveira JT, Mellin C, Lee S, Saber WA, Wang AJ, Ichida JK, Chiu IM, Barrett L, Huebner EA, Bilgin C, Tsujimoto N, Brenneis C, Kapur K, Rubin LL, Eggan K, Woolf CJ. Modeling pain in vitro using nociceptor neurons reprogrammed from fibroblasts. Nat Neurosci 2015;18(1):17–24.

[112] Whittington N, Cunningham D, Le TK, De Maria D, Silva EM. Sox21 regulates the progression of neuronal differentiation in a dose-dependent manner. Dev Biol 2015;397(2):237–247.

[113] Wieskopf JS, Mathur J, Limapichat W, Post MR, Al-Qazzaz M, Sorge RE, Martin LJ, Zaykin DV, Smith SB, Freitas K. The nicotinic α6 subunit gene determines variability in chronic pain sensitivity via cross-inhibition of P2X2/3 receptors. Science translational medicine 2015;7(287):287ra272–287ra272.

[114] Willis DE, Twiss JL. Profiling axonal mRNA transport. Methods Mol Biol 2011;714:335–352.

[115] Wu KY, Hengst U, Cox LJ, Macosko EZ, Jeromin A, Urquhart ER, Jaffrey SR. Local translation of RhoA regulates growth cone collapse. Nature 2005;436(7053):1020–1024.

[116] Wu S, Marie Lutz B, Miao X, Liang L, Mo K, Chang YJ, Du P, Soteropoulos P, Tian B, Kaufman AG, Bekker A, Hu Y, Tao YX. Dorsal root ganglion transcriptome analysis following peripheral nerve injury in mice. Mol Pain 2016;12.

[117] Xie W, Schultz MD, Lister R, Hou Z, Rajagopal N, Ray P, Whitaker JW, Tian S, Hawkins RD, Leung D. Epigenomic analysis of multilineage differentiation of human embryonic stem cells. Cell 2013;153(5):1134–1148.

[118] Yaguchi H, Okumura F, Takahashi H, Kano T, Kameda H, Uchigashima M, Tanaka S, Watanabe M, Sasaki H, Hatakeyama S. TRIM67 protein negatively regulates Ras activity through degradation of 80K-H and induces neuritogenesis. Journal of Biological Chemistry 2012;287(15):12050–12059.

[119] Yin K, Deuis JR, Lewis RJ, Vetter I. Transcriptomic and behavioural characterisation of a mouse model of burn pain identify the cholecystokinin 2 receptor as an analgesic target. Mol Pain 2016;12.

[120] Yoo S, Van Niekerk EA, Merianda TT, Twiss JL. Dynamics of axonal mRNA transport and implications for peripheral nerve regeneration. Experimental neurology 2010;223(1):19–27.

[121] You L, Wu J, Feng Y, Fu Y, Guo Y, Long L, Zhang H, Luan Y, Tian P, Chen L. APASdb: a database describing alternative poly (A) sites and selection of heterogeneous cleavage sites downstream of poly (A) signals. Nucleic acids research 2014;43(D1):D59–D67.

[122] Young-Pearse T, Ivic L, Kriegstein A, Cepko C. Characterization of mice with targeted deletion of glycine receptor alpha 2. Molecular and cellular biology 2006;26(15):5728–5734.

[123] Zaitseva M, Vollenhoven BJ, Rogers PA. In vitro culture significantly alters gene expression profiles and reduces differences between myometrial and fibroid smooth muscle cells. Mol Hum Reprod 2006;12(3):187–207.

